# Shared Neural Signatures of Socioeconomic Status, Scarcity, and Neighborhood Threat in Youth

**DOI:** 10.1101/2025.06.11.659188

**Authors:** K. Brosch, L. Wiersch, E. Christensen, D. Rakesh, J. Hanson, R.M. Schwartz, E. Dhamala

**Author notes:** **Corresponding author**: Dr. K. Brosch.

## Abstract

Early life adversity is a known risk factor for psychopathology, yet the neurodevelopmental impacts of distinct types of adversity remain unclear. Using machine learning, we examined how adversity (physical and sexual abuse, neighborhood threat, scarcity, household dysfunction, prenatal substance exposure, parental psychopathology) and socioeconomic status are associated with cortical thickness in youth. We used data from the Adolescent Brain Cognitive Development Study at three time points: baseline (*N*=6,908, ages 9–10), two-year follow-up (*N*=5,808), and four-year follow-up (*N*=2,245). Cortical thickness was linked to socioeconomic status and neighborhood threat at all time points, and scarcity at baseline. The strongest negative associations were in medial temporal and occipital regions. Our analyses revealed neural effects of socioeconomic status, neighborhood threat, and scarcity, converging on regions involved in memory, visual processing, and higher-order cognition. These findings suggest consistent neural signatures linked to socioeconomic disadvantage, highlighting the importance of addressing inequality to promote neurodevelopmental health.

## Main

Early life adversity encompasses a wide spectrum of pre- and postnatal stressors that occur during sensitive periods and can shape emotional, cognitive, physical and brain development^1–3^. Such early life stressors include prenatal substance exposure, sexual abuse, parental psychopathology, neglect, economic hardship, neighborhood violence, community violence, and housing instability ^1,4–6^. Both chronic exposure to early life adversity and isolated acute incidents have been associated with changes in brain development and elevated risk for mental health problems ^5,7–15^.

Exposure to early life adversities can disrupt brain maturation, stress regulation, and socioemotional development, even in the absence of clinical symptoms^9,16^. Previous research on early life adversity has often relied on small samples, univariate models, or examined only one adversity factor at a time^10,17–19^. This fragmented approach has limited our understanding of how different adversity types may interact or share common neural pathways, making it difficult to capture the full complexity of early life adversity’s impact on development ^20^. There is a growing need for large-scale, data-driven approaches that can capture both shared and distinct neural influences of different forms of early life adversity on adolescent development ^21,22^.

To better characterize adversity, dimensional models have proposed distinctions such as threat, deprivation, and unpredictability ^23,24^. Threat-related adversity includes exposure to violence, sexual or physical abuse, and neighborhood violence, whereas deprivation encompasses reduced cognitive and emotional input, including neglect and resource scarcity ^25^. In earlier studies, these dimensions show distinct neurodevelopmental signatures: threat was associated with smaller cortical surface area in the prefrontal cortex, while deprivation has been linked to larger cortical thickness in the occipital cortex, insula, and cingulate ^26,27^.

Cortical thickness refers to the width of gray matter of the human cortex and is a widely used marker of structural brain development^28–30^. During adolescence, cortical thickness decreases as part of normative brain development, driven by processes such as myelination, synaptic pruning, and pubertal maturation ^31,32^. Notably, smaller cortical thickness has been consistently linked to adversity and psychopathology, underscoring its relevance when studying early life adversity ^33^. While the underlying mechanisms are still being uncovered, evidence from both clinical and non-clinical samples suggests that early adversity is associated with a thinner cortex. Similar patterns across various psychiatric conditions further highlight its role as a transdiagnostic marker of neurodevelopmental risk ^32–34^.

Building on general patterns of adversity, emerging research has begun to characterize how different forms of early adversity may exert distinct effects on brain structure. When examining specific threat-related adversity types, childhood sexual abuse has been associated with enduring alterations, including reduced cortical thickness in the somatosensory cortex regions representing the genitalia, decreased gray matter volume in visual cortices, and increased cerebellar volume ^10,11,18^. Physical abuse has similarly been linked to structural changes in frontolimbic circuits, although findings vary depending on the timing, chronicity, and presence of co-occurring exposures ^35,36^.

Structural brain alterations have also been reported in association with deprivation-related adversity. Neighborhood deprivation, which often includes limited access to resources in a child’s immediate environment has been linked to lower cortical thickness in multiple frontal, parietal and occipital regions, as well as lower overall subcortical volumes ^25,37,38^. In contrast, institutional deprivation, marked by extreme, prolonged lack of caregiving and stimulation, such as in certain orphanage settings, has been associated with reduced gray matter volume in medial prefrontal, inferior frontal, and temporal cortices ^39^. Most research has examined early adversity by focusing on one type of exposure or applying a priori theoretical models. These approaches may overlook the complexity of lived experience, as real-world adversity often cuts across boundaries.

Not all forms of early life adversities map cleanly onto either threat, deprivation, or unpredictability, highlighting limitations of a priori theoretical models. For instance, household dysfunction includes experiences that may include all domains of threat, deprivation, and unpredictability^23,24,26,40^. Similarly, prenatal substance exposure, including exposure to alcohol, cannabis, or other specific substances in utero, represents a biologically embedded adversity that occurs prior to birth and exerts lasting effects despite its fixed timing ^19,41–43^. These exposures are also often correlated with postnatal adversity, compounding developmental risk^42,44^.

Prenatal substance exposure has been linked to widespread neurodevelopmental alterations, including reduced cortical thickness, disrupted gyrification, and atypical maturation patterns in regions critical for language, attention and cognition ^45^. In children and adolescents, prenatal alcohol exposure was associated with smaller cortical thickness and less gyrification in the hippocampus, and smaller surface area in the middle temporal gyrus ^41,43^. These brain alterations were found to mediate the relationship between prenatal alcohol exposure and executive functioning deficits ^41^. Prenatal cannabis exposure has been associated with smaller global gray matter volume, higher risk for internalizing and externalizing symptoms, psychotic-like experiences, sleep problems, and poorer cognitive performance ^19,43^.

The associations between adversity and brain development are shaped by age, intensity, and type of exposure, yet these dimensions are rarely considered together. Different developmental stages involve distinct patterns of brain maturation, and adversity experienced during these windows can have varying impacts on neurodevelopment ^4,31^. Moreover, adversity effects are dose-dependent, with each additional exposure during adolescence raising the likelihood of an adult psychiatric disorder by 52 %, illustrating how chronic or repeated adversity compounds risk ^46^. Together, these findings underscore the need for models that can account for the complexity of adversity’s impact across development ^9,17,47,48^.

Large-scale datasets, such as The Adolescent Brain and Cognitive Development (ABCD) Study, provide an excellent resource to address these complexities. By following over 11,000 children from middle childhood into early adulthood, the study is collecting comprehensive data on mental health, brain development, early life adversity, and environmental influences ^49^. Through its longitudinal assessment, the ABCD Study enables the investigation of how type and severity of adversity, age, and sex influence trajectories of brain development ^49,50^. Data-driven approaches are especially useful in this context, as they allow for the detection of multivariate patterns in highly co-occurring exposures while minimizing reliance on a priori theoretical assumptions ^21^.

In a recent data-driven effort, Orendain and colleagues used exploratory factor analysis across 14 ABCD baseline questionnaires to derive six latent adversity factors, which they labeled “physical and sexual violence”, “prenatal substance exposure”, “scarcity”, “parental psychopathology”, “neighborhood threat”, and “household dysfunction”^6^. Each factor captured a specific domain of adversity. As an example, scarcity reflected material deprivation such as food insecurity and utility shutoffs, while neighborhood threat encompassed perceived danger and crime in the child’s community. These adversity factors varied in prevalence, with some form of parental psychopathology reported in over 80% of participants and physical and sexual violence reported in 7.3% ^6^. All adversity dimensions, except scarcity, were significantly associated with increased internalizing (anxious, depressed, withdrawn) and externalizing (rule-breaking, aggression) symptoms. Moreover, adversity exposure was disproportionately reported among racially and ethnically minoritized youth and those from lower socioeconomic backgrounds, consistent with prior literature on structural inequality and developmental risk ^6,51^.

Together, these findings highlight that adversity rarely occurs in isolation and is often embedded within broader socioeconomic conditions. Further, socioeconomic status is closely tied to many forms of adversity and must be considered when studying early life adversity. It represents both an independent risk factor for altered neurodevelopment and often a proxy for exposure to other adversities^22,52–55^. This multifaceted construct can be captured using indicators such as household income, income-to-needs ratio, and parental education ^5,25,56^. Children from lower socioeconomic backgrounds are disproportionately exposed to early life adversity, contributing to long-term developmental vulnerability ^57^. Socioeconomic status itself has been linked to poorer cognitive and academic performance, greater mental health risk, and altered brain development ^6,25,51,58–61^. These neural differences are especially evident in prefrontal regions and limbic regions involved in emotion and memory, including the amygdala and hippocampus ^60–63^.

Previous work has documented sex-specific differences in the prevalence of certain early life adversities, such as childhood sexual abuse^64^. In addition to differences in prevalence, adversity may also have sex-specific effects on neurodevelopment, underscoring the importance of sex-stratified analyses ^10,65,66^. While some research has focused on specific subgroups, such as young women, large-scale studies directly comparing adversity-related brain effects across sexes remain scarce^10,18,20^.

In this study, we use brain-based predictive models in the ABCD dataset to examine how six dimensions of early life adversity and socioeconomic status (measured using household income) relate to cortical thickness across three timepoints in adolescence (ages 9–14) ^6^. We focus on cortical thickness, as it is a sensitive marker of brain development with robust links to early adversity ^32,33^. We applied multivariate machine learning to capture distributed cortical patterns that univariate approaches may overlook, allowing us to investigate both shared and distinct effects of adversity across time and by sex ^67^. Based on prior work, we expected predominantly negative associations with cortical thickness, distinct patterns for threat- and deprivation-related adversity, and sex-specific effects for physical and sexual abuse ^4,10,18,25,32,33,63,68^. Our findings revealed consistent prediction patterns for neighborhood threat, scarcity, and socioeconomic status across timepoints, suggesting that these exposures may reflect enduring, brain-based signatures of broader socioeconomic disadvantage. By identifying adversity dimensions with stable cortical associations over development, our work highlights neurobiological pathways through which adversity may shape mental and behavioral health and underscores the need to address structural inequality early in life.

## Results

We used separate brain-based predictive models to explore how patterns of cortical thickness are linked to different types of early life adversity and socioeconomic status at three time-points. To do this, we applied a multivariate approach that considers all brain regions simultaneously rather than one at a time. By using brain features to predict adversity exposure, we aimed to capture multivariate (i.e., high-dimensional) associations across the cortex. This allowed us to investigate which patterns across the brain were most strongly associated with each adversity factor. While we use the term “prediction”, these models reflect associations observed at the same point in time and do not imply causality. We first evaluated whether the models could reliably predict adversity scores and then identified the brain regions that contributed most to those predictions.

### Sample Characteristics and Measurement Overview

To ensure data quality, we filtered participants based on data completeness and quality control criteria (see Supplementary Figures 1, 3, and 5). This filtering resulted in a final sample that included a higher proportion of White participants and youth from higher-income households, and a lower proportion of Black and Asian participants compared to those excluded (see Supplementary Table 1). Distributions of adversity measures are shown in Supplementary Figures 2A, 4A, and 6A. Adversity measures showed weak to moderate correlations (Spearman’s *r* = 0.03-0.35, p <.001; Supplementary Figures 2B, 4B, and 6B), indicating that each measure captured a distinct domain of early life adversity. We conducted cross-sectional analyses at three time-points to examine associations between adversity and cortical thickness. As a result, participants may have been included in one, two, or all three time points. Sample overlap across time-points is visualized in Supplementary Figure 7. A total of 1,275 participants were included at all three time points.

### At baseline, cortical thickness significantly predicts neighborhood threat, scarcity, and socioeconomic status

At baseline, data were available for seven factors: the six adversity factors (physical abuse and sexual abuse, neighborhood threat, scarcity, household dysfunction, prenatal substance exposure, parental psychopathology), and socioeconomic status. Cortical thickness was significantly associated with neighborhood threat (prediction accuracy, *r*=0.141, *p_FDR_*<0.001), scarcity (*r*=0.131, *p_FDR_*<0.001), and socioeconomic status (*r*=0.274, *p_FDR_*<0.001). Cortical thickness was not significantly associated with physical and sexual abuse (*r*=-0.004, *p_FDR_*=0.550), prenatal substance exposure (*r*=0.028, *p_FDR_*=0.200), household dysfunction (*r*=0.028, *p_FDR_*=0.327), or parental psychopathology (*r*=0.046, *p_FDR_*=0.067). Prediction accuracies for all models and time-points are visualized in Figure 1. Feature weights are visualized for the significant models in Figure 2.

**Figure 1:**
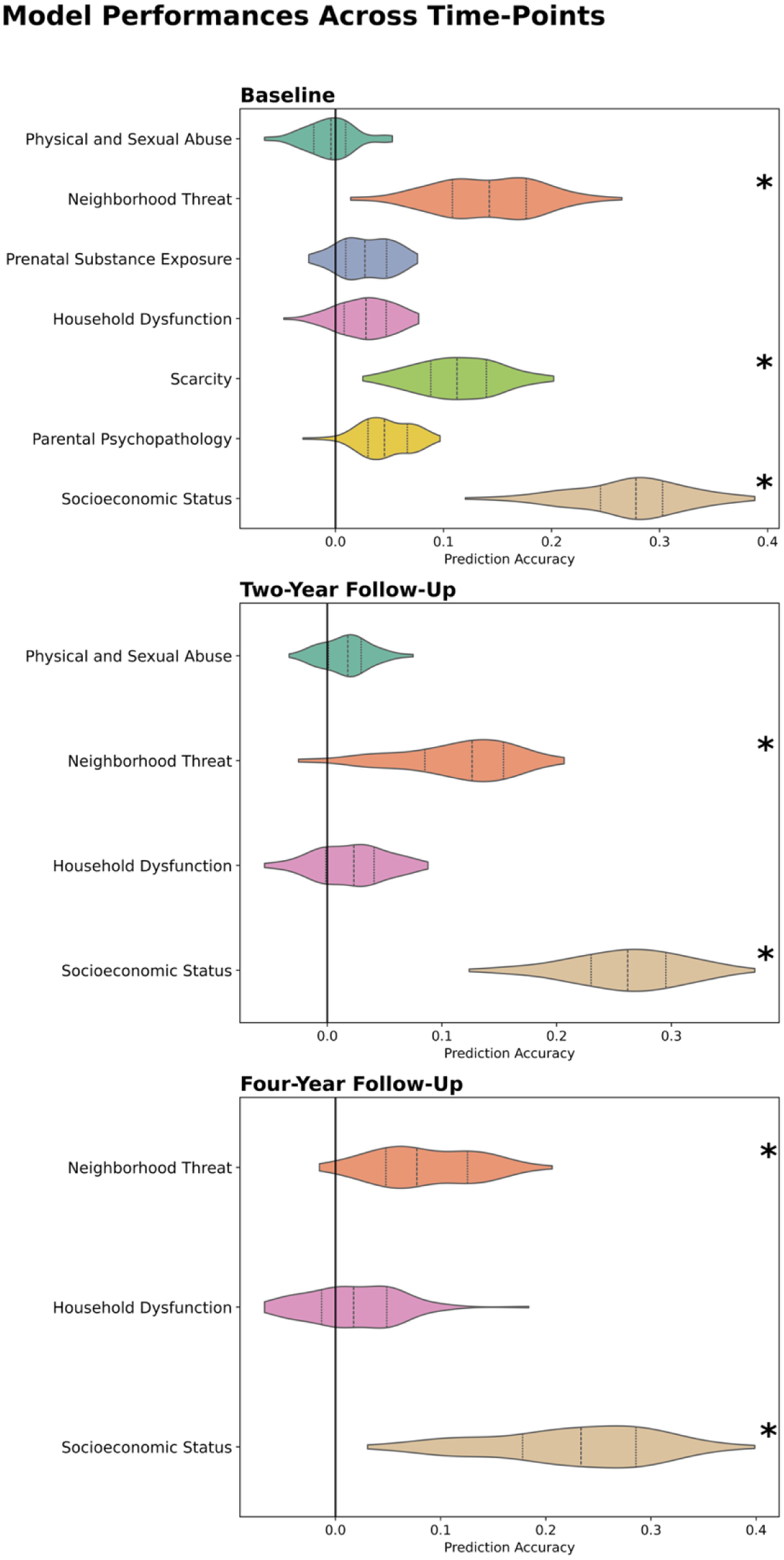
Model prediction accuracies (correlation coefficient between observed and predicted scores) for models trained to predict cortical thickness at the three time-points, top panel: baseline, middle panel: two-year follow-up, bottom panel: four-year follow-up. Asterisks indicate significant predictive models after FDR-error correction. The shape of the violin plots indicates the distribution of values, the dashed lines indicate the median, and the dotted lines indicate the interquartile range.

**Figure 2:**
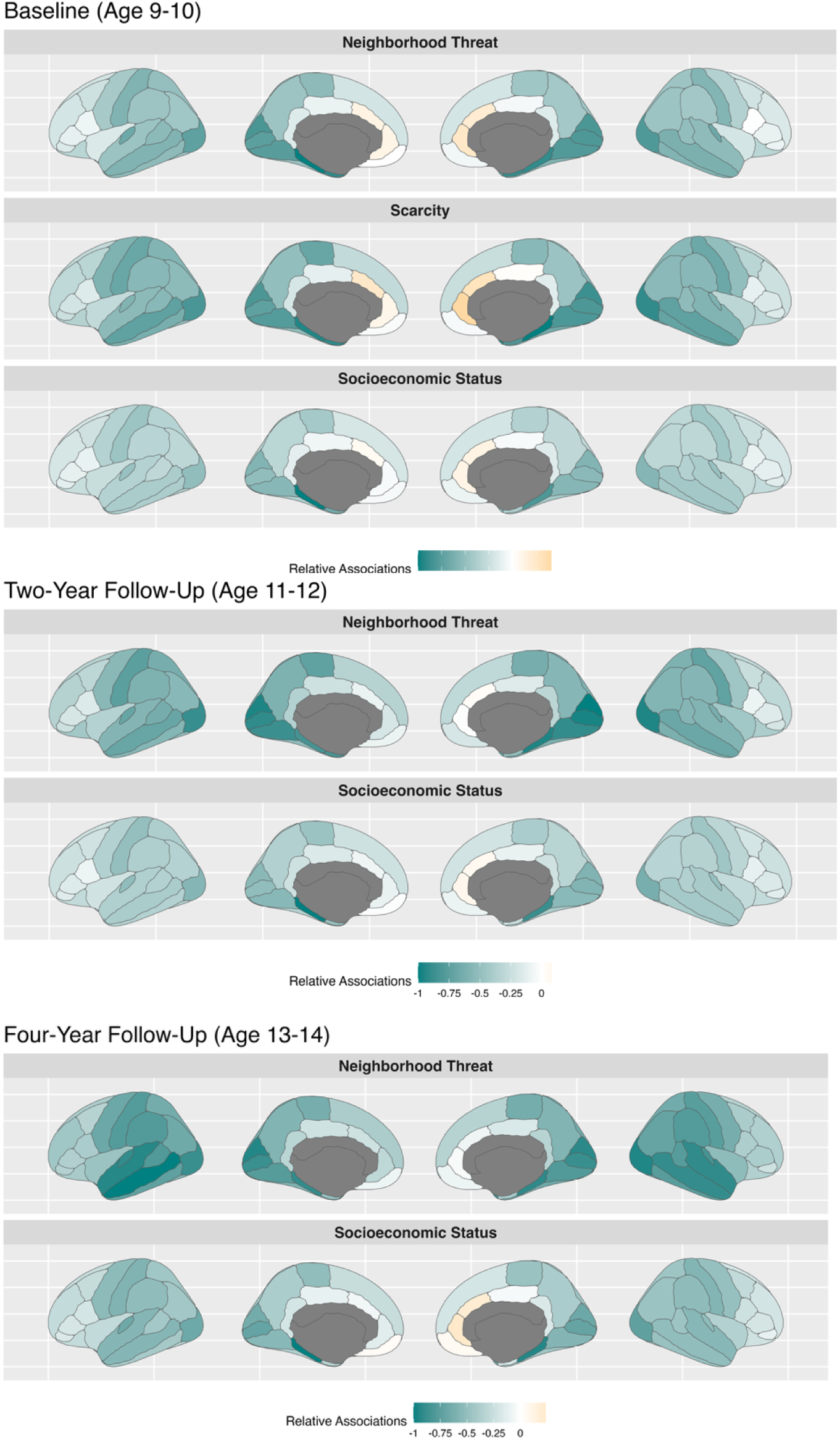
Relative associations between cortical thickness and significant factors at the three time-points. Deeper green colors indicate a stronger negative association, deeper yellow colors indicate a stronger positive association. To facilitate visualization, association values were divided by the maximum value for that model. Lateral (outer) and medial (inner) surfaces for left (left) and right (right) hemispheres are shown.

### At two-year follow-up, cortical thickness significantly predicts neighborhood threat and socioeconomic status

At two-year follow-up, data were available for four factors: physical and sexual abuse, neighborhood threat, household dysfunction, and socioeconomic status. Cortical thickness was significantly associated with neighborhood threat (*r*=0.115, *p_FDR_*=0.002) and socioeconomic status (*r*=0.261, *p_FDR_*<0.001), but not with physical and sexual abuse (*r*=0.017, *p_FDR_*=0.264) or household dysfunction (*r*=0.021, *p_FDR_*=0.253).

### At four-year follow-up, cortical thickness significantly predicts neighborhood threat and socioeconomic status

At four-year follow-up, data on neighborhood threat, household dysfunction and socioeconomic status were available. Cortical thickness was significantly associated with neighborhood threat (*r*=0.085, *p_FDR_*=0.036) and socioeconomic status (*r*=0.223, *p_FDR_*<0.001). There was no significant association with household dysfunction (*r*=0.017, *p_FDR_*=0.636).

### Brain-Adversity Associations reveal few sex-specific findings

We conducted separate sex-specific analyses for boys and girls, which revealed similar findings and feature weights across the three time points compared to the combined sample (see Supplementary Figures 8-10). Notably, at baseline, some regions showed descriptively higher feature weight scores in girls compared to boys, despite the overall similarity in patterns. While our results revealed few sex-specific differences, conducting sex-stratified analyses remains essential given known sex differences in the prevalence and impact of certain adversity types, such as childhood sexual abuse. Such analyses can uncover nuanced effects even when overall patterns appear similar.

Overall, these findings suggest that, within the available data, socioeconomic status and neighborhood threat are consistently associated with cortical thickness across all three time-points.

We then Haufe-transformed the feature weights to increase reliability and interpretability and examined the specific associations captured by each model ^69,70^. Here, we focus our results on models that yielded significant results. Unlike traditional associations, feature weights reflect how much each region contributed to the model’s ability to predict adversity or socioeconomic status. The figures below visualize this relative importance of each brain region based on its contribution to the individual predictions.

### Neighborhood threat, scarcity, and socioeconomic status influence overlapping brain regions

<u>At baseline</u>, neighborhood threat, scarcity, and socioeconomic status mapped onto overlapping regions, which were predominantly the bilateral medial temporal and occipital regions. We detected mostly negative associations, indicating smaller cortical regional thickness was associated with higher adversity exposure. Here, we detected the strongest associations for all three measures in the bilateral parahippocampal and entorhinal cortices, bilateral cunei, bilateral lingual gyri, bilateral pericalcarine and lateral occipital cortices, and bilateral temporal poles. We identified positive associations (indicating greater cortical thickness was associated with higher adversity exposure) for all three measures in the bilateral rostral and caudal anterior cingulate, and bilateral frontal poles (Figure 2).

<u>At two-year follow-up,</u> neighborhood threat and socioeconomic status were associated with cortical thickness in similar regions, which were in turn similar to the associations observed at baseline. The strongest negative associations were in the bilateral parahippocampal and entorhinal cortices, bilateral cunei, bilateral lingual gyri, bilateral pericalcarine and lateral occipital cortices, and bilateral temporal poles. Positive associations were observed in the bilateral rostral and caudal anterior cingulate, and bilateral frontal poles (Figure 2).

<u>At four-year follow-up,</u> associations between cortical thickness and socioeconomic status were very similar to those observed at both baseline and two-year follow-up. The strongest negative associations were in the bilateral parahippocampal cortices, bilateral temporal poles, bilateral lateral occipital and pericalcarine gyri, right cuneus and left transverse temporal. Positive associations were found in the right rostral and caudal anterior cingulate cortex and right frontal pole (Figure 2).

### Distinct adversity factors and socioeconomic status map onto shared brain regions within and across time-points

To assess whether significant models predicting different factors implicated similar brain regions, we calculated cosine similarities between feature weight vectors, allowing us to evaluate the spatial similarity of brain–exposure associations across models. Cosine similarity values closer to 1 indicate a greater overlap in the spatial distribution of feature weights, suggesting that the models rely on similar brain regions to predict different exposures.

At baseline, neighborhood threat, scarcity, and socioeconomic status were associated with cortical thickness in overlapping brain regions (cosine similarity, s=0.98-0.99), indicating nearly perfect overlap. At two-year follow-up, socioeconomic status and neighborhood thread were associated with cortical thickness in overlapping brain regions (s=0.98). At four-year follow-up, socioeconomic status and neighborhood threat were again associated with cortical thickness in overlapping brain regions (s=0.98), see Supplementary Figure 8.

To investigate if these associations were stable across different time-points, we also calculated cosine similarities for the models that captured significant associations across baseline, two-year, and four-year follow-up (see Supplement). Cosine similarities across time-points and models were consistently very high (s > 0.96), indicating that associations between cortical thickness and neighborhood threat, scarcity and socioeconomic status remained stable over those time points, see Supplementary Figure 11.

### Findings remain robust using income-to-needs ratio and non-overlapping samples

To test whether our results for socioeconomic status were robust when using a more sensitive measure than household income, we re-ran the analyses using the income-to-needs ratio, which compares a family’s income to the poverty threshold, accounting for household size. At baseline, income-to-needs ratio was significantly associated with cortical thickness (*r*=0.220, p_FDR_<0.001), showing similar feature weights as socioeconomic status, Supplementary Figure 12 This indicates that socioeconomic status is a relevant, stable factor and alternate measures support our findings.

To assess whether participant overlap across time points influenced our findings, we re-ran the baseline and 2-year analyses using non-overlapping participant samples, while keeping the 4-year sample unchanged (*n* = 2,245) to retain the largest possible sample at that time point. Specifically, we selected the largest available subsamples at baseline (*n* = 2,413) and 2-year follow-up (*n* = 4,086) with no participant overlap across time points Supplementary Figure 13. These analyses yielded results consistent with the main findings (Supplementary Figure 14), reinforcing the robustness of our results.

Overall, these results indicate that socioeconomic status, neighborhood threat, and scarcity share an overlapping neural signature that is stable across three time-points.

## Discussion

This study leveraged the scale and depth of the ABCD Study to investigate how early life adversity and socioeconomic status relate to cortical thickness across youth. Using predictive modeling in a large sample, we found that socioeconomic status, neighborhood threat, and (at baseline) scarcity were consistently associated with smaller cortical thickness. These three distinct factors showed highly overlapping associations, suggesting that together, they may reflect a common neural signature of broader structural disadvantage. Although fewer adversity factors were available at later time points, the spatial patterns of these associations remained stable, pointing to persistent cortical correlates of socioeconomic adversity across development.

We observed mostly negative associations between adversity and cortical thickness, consistent with our hypotheses and with prior literature. These relationships were strongest in bilateral medial temporal, occipital, and temporal pole regions, with smaller clusters of positive associations in the bilateral rostral and anterior cingulate and frontal pole. These regions are broadly involved in memory, emotion regulation, language, and visual processing, highlighting the potential impact of socioeconomic adversity on multiple cognitive and affective systems ^71–73^.

In contrast, the small number of positive associations in frontal regions may reflect compensatory neural adaptations or individual variability in developmental timing. Together, these patterns point to a distributed and functionally meaningful neural signature of socioeconomic disadvantage that is detectable across multiple stages of adolescence. They also support the idea that early adversity is biologically embedded in brain development, shaping structural features that are foundational for cognition and emotion.

Our findings of smaller cortical thickness associated with socioeconomic status, neighborhood threat, and scarcity align with prior work showing smaller cortical thickness in youth exposed to adversity or disadvantage ^32,42,74^. However, these results require careful interpretation, as multiple mechanisms may underlie these associations. One prominent explanation is accelerated maturation, a process by which adversity exposure may speed up cortical development as an adaptation to challenging environments. A more mature cortex may offer short-term functional advantages by helping children navigate high-stress or resource-limited environments through enhanced cognitive and behavioral efficiency ^9,75^.

Supporting this interpretation, past research has documented accelerated cortical maturation and accelerated epigenetic aging in children from lower socioeconomic backgrounds ^76,77^. However, smaller cortical thickness at the three time-points may not reflect faster versions of typical development, but rather indicate fundamentally different neurodevelopmental pathways. The detected patterns could reflect pre-existing structural differences that emerge early and persist over time, rather than altered developmental timing ^25^. Our analyses cannot distinguish between these competing explanations. Disentangling them will require longitudinal designs that track individual-level developmental trajectories in relation to adversity exposure.

Importantly, regardless of whether these structural differences reflect accelerated maturation or persistent alterations, they are consistently linked to adverse functional outcomes. A broad literature connects socioeconomic disadvantage to poorer mental health, lower academic performance, and greater functional impairments, suggesting that any neurodevelopmental “adaptations” reflected in cortical thickness are insufficient to mitigate the negative long-term impacts of socioeconomic disadvantage ^59–61,78–80^. Further, children from lower socioeconomic backgrounds are more likely to be exposed to additional adversities, leading to greater risk for mental health problems, a pattern described as “double-disadvantage”^66,81,82^. While accelerated maturation may be adaptive in the short term, it may come at a cost, including reduced plasticity, earlier closure of sensitive periods, and long-term trade-offs in flexibility and resilience ^9^.

Certain limitations should be noted. In this study, we used caregiver-reported household income as a proxy for socioeconomic status. This measure complements other metrics used in the literature, such as income-to-needs ratio, parental education, or geocoded indices like the Area Deprivation Index, each capturing distinct aspects of socioeconomic context ^5,68,83,84^. Despite its simplicity, household income aligned with more complex indices such as neighborhood threat and scarcity. Findings remained consistent when using income-to-needs ratio, highlighting the robustness of socioeconomic status as a predictor. Although socioeconomic status, neighborhood threat, and scarcity were only weakly to moderately correlated behaviorally, their neural correlates overlapped, suggesting shared neurobiological pathways. While not interchangeable with more comprehensive measures, household income may still offer a useful, albeit limited, proxy for adversity in settings where more detailed measures are unavailable.

Interestingly, our multivariate results did not replicate previously described univariate associations between cortical thickness and certain adversity factors such as physical and sexual violence, prenatal substance exposure, parental psychopathology, and household dysfunction ^10,13,17–19,41,43^. These discrepancies may be due to differences in analytic strategy, as our multivariate approach may capture different relationships than traditional univariate analyses. Further, differences in sample age, exposure intensity, and how adversity factors were defined and captured across studies may impact findings ^17^. For instance, some studies rely on dichotomous classifications of adversity, which may limit sensitivity to variation in exposure and reduce comparability across analytic frameworks ^17^. In contrast, we treated adversity dimensions as continuous to capture gradations in exposure and increase model sensitivity.

Another consideration for the interpretation of our findings is that we did not observe meaningful changes in the neural correlates of adversity over time. Associations were largely stable across time points, and adversity exposures themselves showed little change across the study period (Supplementary Figure 15). This may reflect limited variability or intensity of exposure in this age range or specific sample. Additionally, some adversity factors were not reassessed at follow-up, limiting our ability to examine change. Selective attrition may also have influenced results, as higher-risk participants or those with lower socioeconomic status may have been more likely to drop out or be excluded due to poor imaging data ^85^. Supporting this, we found lower prevalences of adversity exposure in our sample compared to Orendain et al.’s ABCD baseline sample (e.g., 3.5% vs. 7% for sexual abuse, 81.4% vs. 50.3% for parental psychopathology) ^6^. Our exclusion of siblings and those with missing or poor-quality MRI data may have further reduced the representation of higher-risk groups, which remains a broader challenge in imaging research ^86–88^. Our sample also skewed toward higher-income families (see Supplement Table 1). Finally, some forms of adversity may be underreported, especially interpersonal trauma, which caregivers may be less aware of or less likely to disclose.

Although this study focused specifically on cortical thickness as one marker of brain development, early life adversity is known to affect a broader range of neural systems. Prior work has demonstrated associations between adversity and functional connectivity, particularly in networks supporting emotion regulation and executive functioning ^22,89–94^. These findings suggest that adversity may exert diffuse effects across structural and functional domains ^58,89^. In this study, we did not include race as a covariate, as it often functions as a proxy for socioeconomic status and structural inequality ^12,95^ and because previous research has shown that race-related differences in brain structure can be attributed to differences in exposure to adversity^12^.

Despite these considerations, our findings were robust as we replicated our findings using completely non-overlapping participant samples at the three time-points. This suggests that our results are not driven by repeated measurements of the same individuals. In addition, we replicated our findings for household income (socioeconomic status) using the income-to-needs ratio, a more sensitive measure of socioeconomic status^38^. The convergence of findings across different samples and socioeconomic status measures underscores the reliability of these associations.

Our findings demonstrate the significant and pervasive negative impact of socioeconomic disadvantage on adolescent brain development. Although certain protective factors may buffer some of these effects, the overall burden of disadvantage remains profound ^96–99^. The effects of socioeconomic disadvantage permeate nearly every aspect of a child’s lived experience, including reduced access to high-quality healthcare, nutritious food, and stable, safe housing, as well as increased exposure to environmental hazards, overcrowded living conditions, and neighborhood violence. These disadvantages often extend to diminished educational resources, limited extracurricular opportunities, and lower availability of community support systems ^98,100,101^. Financial strain may also reduce caregiver availability and increase family stress, further compounding risk. Together, these structural and contextual burdens create a chronic, multifaceted environment that may undermine healthy brain development^53,54^. While it is essential to acknowledge these risks, strength-based approaches that highlight resilience and adaptive potential in the face of adversity are equally important for fostering a more comprehensive understanding of developmental outcomes ^66,99^.

We provide a valuable foundation for future work by using multivariate brain-based predictive modeling in a large dataset. By examining six dimensions of early adversity and socioeconomic status across three timepoints, we identified robust, shared cortical thickness patterns, particularly for exposures to neighborhood threat, scarcity, and low socioeconomic status. These findings suggest that some adversity-related alterations may reflect stable neurodevelopmental signatures. While we cannot determine whether these reflect accelerated or blunted maturation, our work offers a critical step toward clarifying these mechanisms. With this multi-timepoint, data-driven framework in place, future studies can investigate whether cortical differences represent altered developmental timing or persistent structural differences ^25^.

Moving forward, future work should apply longitudinal analyses and multimodal data, including structural, epigenetic, and environmental measures. Such approaches will determine whether adversity-related cortical changes reflect altered developmental timing or permanent structural shifts. This knowledge will guide when, how, and for whom to intervene to mitigate early-adversity impacts and inform programs that build resilience, particularly among socioeconomically disadvantaged youth.

## Methods

### Participants

Data from the Adolescent Brain Cognitive Development (ABCD) study (ABCD 5.1 release) were used to establish associations between adverse childhood experiences and cortical thickness at three time points. The ABCD study assesses the development of over 11,000 children (aged 9-10 at baseline) from 21 sites across the United States. Participants complete extensive assessments, including neuroimaging scans, cognitive assessments, and mental health questionnaires, among others^49,102^. Inclusion criteria for the ABCD study can be found here^49^. In our study, we included individuals with complete and usable structural imaging data, socioeconomic status data, and complete data for all items necessary to calculate the adversity factors at the specific time point. We excluded intersex individuals, individuals with clinically relevant brain structural variations, individuals with T1 data not recommended for inclusion. We also excluded one or more members from sibling pairs to ensure no siblings were included in the sample, and excluded sites with fewer than 20 participants. At baseline, we included 6,908 individuals (3,282 females), at two-year follow-up we included 5,808 individuals (2,686 females), and at four-year follow-up we included 2,245 individuals (1,044 females).

### Measures

#### Early life adversity

We use the term early life adversity to refer to a range of stressors experienced prenatally and during youth. We considered six adversity factors, previously identified using factor analysis in the ABCD study by Orendain and colleagues^6^: 1) Physical and Sexual Violence, 2) Neighborhood Threat, 3) Prenatal Substance Exposure, 4) Household Dysfunction, 5) Scarcity, and 6) Parental Psychopathology. We calculated continuous scores for each factor from 47 items using 14 different questionnaires assessed in the ABCD study, as described by Orendain et al., see Supplement. All items were rated by the parent or caregiver, except for the household dysfunction items, which were rated by the youth.

We additionally included socioeconomic status, operationalized as household income, as we expected neighborhood threat and scarcity to correlate with income. Not all questionnaires necessary to calculate the specific adversity factors were collected at all timepoints. At baseline, data for all six adversity factors were available. At two-year follow-up, only physical and sexual abuse, neighborhood threat, and household dysfunction data were available. At four-year follow-up, only neighborhood threat and household dysfunction data were available. Socioeconomic status was assessed at all three time-points. The available scores for each of the time-points were calculated as described by Orendain^6^; a detailed description can be found in the Supplemental Material. Socioeconomic status was assessed on a 10-point Likert scale capturing income brackets from <$5,000 to >$200,000. We reverse-coded the scale so that higher scores reflected greater disadvantage, allowing for direct comparison with the adversity factors (e.g., 1 = >$200,000, 2 = $100,000 - $199,999). We used Spearman correlations to investigate associations between adversity factors. To explore the robustness of our findings, we additionally conducted brain-based predictive analyses using the income-to-needs ratio, a more sensitive socioeconomic measure that accounts for household size and individual poverty thresholds. Specifically, we calculated this ratio by taking the median value of each income bracket and dividing it by the federal poverty threshold corresponding to the respective household size ^38^.

#### Neuroimaging Data

T1-weighted images were collected using 3T MRIs. The ABCD neuroimaging and (pre)processing protocols can be found here ^102,103^. We extracted mean cortical thickness values from 34 bilateral cortical regions defined by the Desikan-Killiany atlas, as part of the ABCD study’s standard FreeSurfer preprocessing pipeline.

### Statistical Analyses

#### Predictive Modeling

We applied a cross-validated brain-based predictive modeling framework using linear ridge regression algorithms^104–109^. This framework minimizes overfitting, prevents data leakage, and captures robust, interpretable associations between brain measures and behavioral variables.

In this study, we used these models to predict adversity factors and socioeconomic status from regional cortical thickness across three time-points (7 factors at baseline, four factors at two-year follow-up, and three factors at four-year follow-up). For each time point and each factor, we trained models using 68 cortical thickness features from the Desikan-Killiany atlas. We first randomly split the dataset into 100 distinct training and test sets, ensuring that participants from the same data collection site were not split across sets. Within each training set, we optimized the regularization hyperparameter using three-fold cross-validation, keeping data from sites intact within folds.

After optimization, we trained the model on the full training set and evaluated performance on the corresponding test set. Model performance was evaluated based on prediction accuracy, defined as the correlation between predicted and observed values, in line with prior work ^70,86,104,110^. We assessed within-factor prediction accuracy (how accurately each factor was predicted from cortical thickness at a given time point). We further tested whether model performance exceeded chance using permutation-based null models (generated using 1000 repetitions). P-values were calculated as the proportion of null models with prediction accuracy equal to or greater than the observed value.

To correct for multiple comparisons, we applied the Benjamini-Hochberg procedure to control the false discovery rate (FDR). At baseline, data to calculate all six adversity factors and socioeconomic status were available. At two-year follow-up, data to calculate physical and sexual abuse, neighborhood threat, household dysfunction, and data on socioeconomic status were available. At four-year follow-up, data to calculate neighborhood threat, household dysfunction, and data on socioeconomic status were available. We corrected p-values for 7 tests at baseline, 4 tests at two-year follow-up, and 3 tests at four-year follow-up based on the number of adversity factors available at each time point. Results were considered statistically significant at a threshold of *p*_FDR_= 0.050. Finally, we conducted sex-specific analyses (see Supplement), examining associations between adversity factors and cortical thickness separately for boys and girls at each time point.

#### Covariates

In line with methodological considerations in our previous work, we did not include covariates such as age and sex, as these may mask important findings in the brain-based models^104,111,112^. Covariate adjustment assumes a linear and independent association between outcome and covariate, which may not accurately reflect complex biology. Excluding covariates enabled us to detect more robust brain-behavior associations as the models can reflect a larger range of variation in the neuroimaging data. Importantly, the ABCD sample varies only narrowly in age at each time point, with a standard deviation of only 7 months at each assessment, which reduces the likelihood that age-related differences drive our findings. Additionally, we conducted sex-specific analyses to explore potential differences in associations by sex. We did not use intracranial volume as a covariate, in line with recommendations for cortical thickness analyses^109,113^.

#### Feature Weights

To increase interpretability and reliability, we used the Haufe transformation to transform the obtained feature weights from the predictive models ^69^. We then calculated mean feature importance for each set of the models. To evaluate the similarity between regional features associated with the adversity factors, we calculated cosine similarities, which is calculated using the dot product of two vectors divided by the product of their magnitudes. Values range from −1 to 1, with 1 indicating complete similarity, −1 indicating complete dissimilarity, and 0 indicating orthogonality.

#### Software and Code Availability

All analyses were conducted in Python (version 3.12.3) using Jupyter Notebook and RStudio (4.3.3). Feature weights were visualized using RStudio, and prediction accuracy plots were generated using the Seaborn package in Python. All code used to generate the results will be made available on GitHub.

## CRediT authorship contribution statement

**KB**: Conceptualization, Writing - original draft, Software, Formal Analysis, Visualization; **LW**: Software, Formal Analysis, Writing - review & editing**; EC**: Writing - review & editing; **DR:** Writing - review & editing, **JH:** Writing - review & editing, **RS:** Writing - review & editing **, ED**: Methodology, Software, Writing - review & editing, Supervision, Funding acquisition.

## Acknowledgements

During the preparation of this work the authors used LLM to improve readability and language. After using this tool/service, the authors reviewed and edited the content as needed and take full responsibility for the content of the publication.

