## Supplementary Materials for "Shared Neural Signatures of Socioeconomic Status, Scarcity, and Neighborhood Threat in Youth"

Brosch *et al.*

**Supplementary Table 1: Demographic information.**

Demographic information (age, race/ethnicity, and socioeconomic status/income) for all subjects from the ABCD 5.1. Data Release. Demographic information is reported separately for subjects who were included and excluded from analyses at A) Baseline, B) 2-year follow-up, and C) 4-year follow-up, based on the criteria described in the Supplemental Methods. Reported proportions (%) may not sum to exactly 100% due to rounding.

**A) Baseline, *N* = 11,868**

|  | **Included** (*n*=6,908) | **Excluded** (*n*=4,960) |
| --- | --- | --- |
| **Sex (Count, proportion)** |  |  |
| *Male* | 3,626 (52.49%) | 2,562 (51.65%) |
| *Female* | 3,282 (47.51%) | 2,395 (48.29%) |
| *Intersex-Male* | 0 (0.0%) | 3 (0.06%) |
| **Age in months (mean, SD)** | 118.85 (7.41) | 119.16 (7.61) |
| **Race/Ethnicity (count, %)** |  |  |
| *White* | 3788 (54.83%) | 2385 (48.08%) |
| *Black* | 894 (12.94%) | 890 (17.94%) |
| *Hispanic* | 1390 (20.12%) | 1020 (20.56%) |
| *Asian* | 134 (1.94%) | 118 (2.38%) |
| *Other* | 702 (10.16%) | 545 (10.99%) |
| *NA* | 0 (0.0%) | 2 (0.04%) |
| **Income Category (count, %)** |  |  |
| *Less than $5,000* | 260 (3.76%) | 157 (3.17%) |
| *$5,000 - $11,999* | 276 (4.0%) | 145 (2.92%) |
| *$12,000 - $15,999* | 174 (2.52%) | 99 (2.0%) |
| *$16,000 - $24,999* | 321 (4.65%) | 202 (4.07%) |
| *$25,000 - $34,999* | 406 (5.88%) | 248 (5.0%) |
| *$35,000 - $49,999* | 566 (8.19%) | 368 (7.42%) |
| *$50,000 - $74,999* | 931 (13.48%) | 567 (11.43%) |
| *$75,000 - $99,999* | 1,001 (14.49%) | 569 (11.47%) |
| *$100,000 - $199,999* | 2,139 (30.96%) | 1172 (23.63%) |
| *≥ $200,000* | 834 (12.07%) | 416 (8.39%) |
| *Refuse to answer* | 0 (0.0%) | 511 (10.3%) |
| *Don't know* | 0 (0.0%) | 504 (10.16%) |
| *NA* | 0 (0.0%) | 2 (0.04%) |

**B) 2-year Follow-up, *N* = 10,908**

|  | **Included** (*n*=5,808) | **Excluded** (*n*=5,100) |
| --- | --- | --- |
| **Sex (Count, proportion)** |  |  |
| *Male* | 3,122 (53.75%) | 2,602 (51.02%) |
| *Female* | 2,686 (46.25%) | 2,495 (48.92%) |
| *Intersex-Male* | 0 (0.0%) | 3 (0.06%) |
| **Age in months (mean, SD)** | 143.28 (7.76) | 145.46 (8.17) |
| **Race/Ethnicity (count, %)** |  |  |
| *White* | 3257 (56.08%) | 2578 (50.55%) |
| *Black* | 713 (12.28%) | 836 (16.39%) |
| *Hispanic* | 1130 (19.46%) | 1018 (19.96%) |
| *Asian* | 114 (1.96%) | 115 (2.25%) |
| *Other* | 594 (10.23%) | 552 (10.82%) |
| *NA* | 0 (0.0%) | 1 (0.02%) |
| **Income Category (count, %)** |  |  |
| *Less than $5,000* | 174 (3.0%) | 149 (2.92%) |
| *$5,000 - $11,999* | 177 (3.05%) | 124 (2.43%) |
| *$12,000 - $15,999* | 120 (2.07%) | 97 (1.9%) |
| *$16,000 - $24,999* | 202 (3.48%) | 167 (3.27%) |
| *$25,000 - $34,999* | 330 (5.68%) | 216 (4.24%) |
| *$35,000 - $49,999* | 443 (7.63%) | 288 (5.65%) |
| *$50,000 - $74,999* | 802 (13.81%) | 512 (10.04%) |
| *$75,000 - $99,999* | 820 (14.12%) | 554 (10.86%) |
| *$100,000 - $199,999* | 1,953 (33.63%) | 1,438 (28.2%) |
| *≥ $200,000* | 787 (13.55%) | 656 (12.86%) |
| *Refuse to answer* | 0 (0.0%) | 464 (9.1%) |
| *Don't know* | 0 (0.0%) | 434 (8.51%) |
| *NA* | 0 (0.0%) | 1 (0.02%) |

**C) 4-year Follow-up, *N* = 4,688**

|  | **Included** (*n*=2,245) | **Excluded** (*n*=2,443) |
| --- | --- | --- |
| **Sex (Count, proportion)** |  |  |
| *Male* | 1201 (53.5%) | 1255 (51.37%) |
| *Female* | 1044 (46.5%) | 1187 (48.59%) |
| *Intersex-Male* | 0 (0.0%) | 1 (0.04%) |
| **Age in months (mean, SD)** | 168.52 (8.22) | 169.37 (8.11) |
| **Race/Ethnicity (count, %)** |  |  |
| *White* | 1293 (57.59%) | 1364 (55.83%) |
| *Black* | 219 (9.76%) | 283 (11.58%) |
| *Hispanic* | 447 (19.91%) | 509 (20.84%) |
| *Asian* | 53 (2.36%) | 55 (2.25%) |
| *Other* | 233 (10.38%) | 232 (9.5%) |
| *NA* | 0 (0.0%) | 0 (0.0%) |
| **Income Category (count, %)** |  |  |
| *Less than $5,000* | 46 (2.05%) | 35 (1.43%) |
| *$5,000 - $11,999* | 46 (2.05%) | 40 (1.64%) |
| *$12,000 - $15,999* | 38 (1.69%) | 26 (1.06%) |
| *$16,000 - $24,999* | 93 (4.14%) | 71 (2.91%) |
| *$25,000 - $34,999* | 114 (5.08%) | 84 (3.44%) |
| *$35,000 - $49,999* | 161 (7.17%) | 142 (5.81%) |
| *$50,000 - $74,999* | 272 (12.12%) | 228 (9.33%) |
| *$75,000 - $99,999* | 320 (14.25%) | 269 (11.01%) |
| *$100,000 - $199,999* | 760 (33.85%) | 759 (31.07%) |
| *≥ $200,000* | 395 (17.59%) | 384 (15.72%) |
| *Refuse to answer* | 0 (0.0%) | 243 (9.95%) |
| *Don't know* | 0 (0.0%) | 160 (6.55%) |
| *NA* | 0 (0.0%) | 2 (0.08%) |

Note: Household income was assessed on a 10-point scale by the ABCD study (reversed here for interpretability), with 1 indicating the highest bracket (≥ $200,000) and 10 the lowest (< $5,000). Most participants reported incomes in the $100,000–$199,999 range (mode = 2), and over 40% of families reported household incomes above $100,000. The median and mean values both fell between $75,000 and $99,999. In contrast, the U.S. Census Bureau reports a median household income of approximately $84,000 among families with children.

**Participant Inclusion Pipeline at Baseline**

*n* = 11,868

*n* = 3 excluded: intersex individuals

n = 11,865

*n* = 140 excluded: missing brain structural data

*n* = 11,725

*n* = 11,267

*n* = 458 excluded: T1 data not recommended for inclusion, (“imgincl_t1w_include = 0”)

*n* = 415 excluded: T1 data incidental findings reported, (“mrif_score = 3 or 4”)

*n* = 10,852

*n* = 8,216

*n* = 1,289 excluded: siblings, (duplicate “rel_family_id”)

*n* = 6,927

***n* = 6,908**

*n* = 19 excluded: site22 < 20 individuals, (“site_id < 20”)

*n* = 2,636 excluded: missing behavioral data to calculate adversity factors or income

**Supplementary Figure 1: Participant inclusion and exclusion workflow at baseline.**

Overview of the process used to include participants in the analyses at baseline.

A. **Distribution of Behavioral Scores at Baseline**

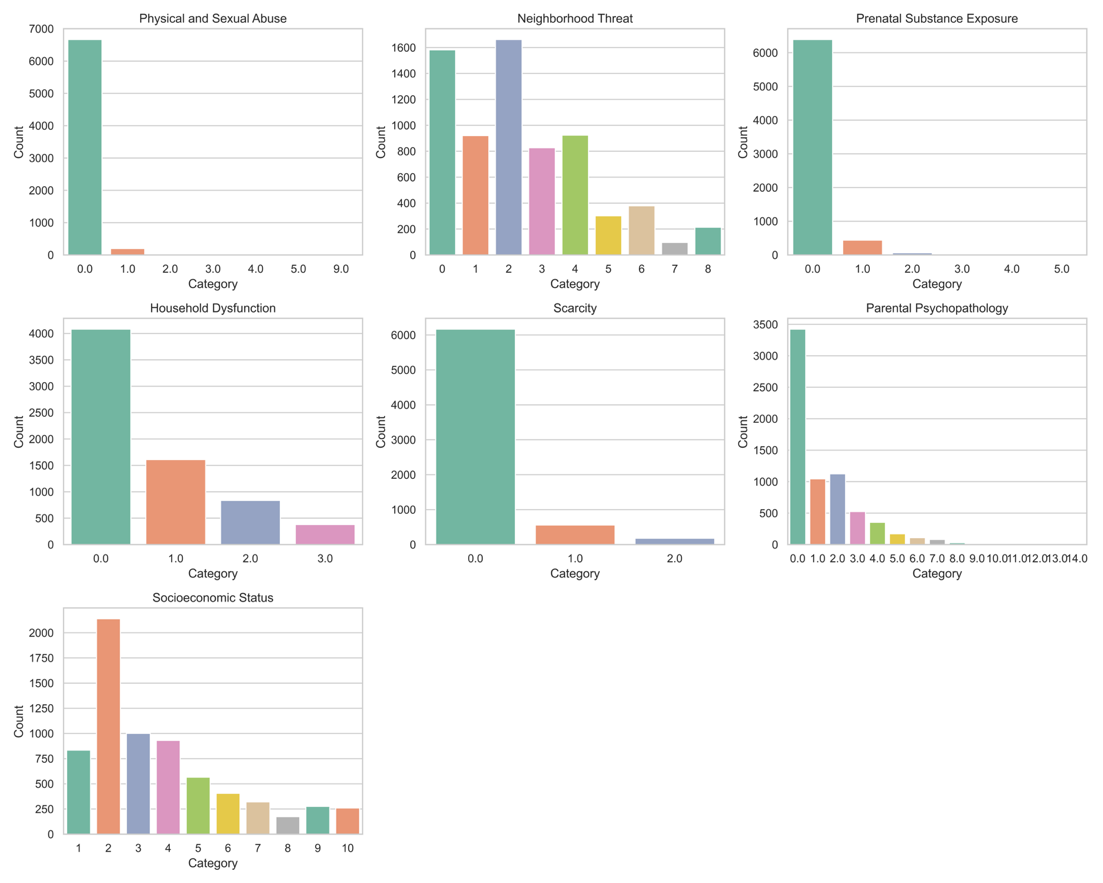

B. **Correlation of Behavioral Scores at Baseline**

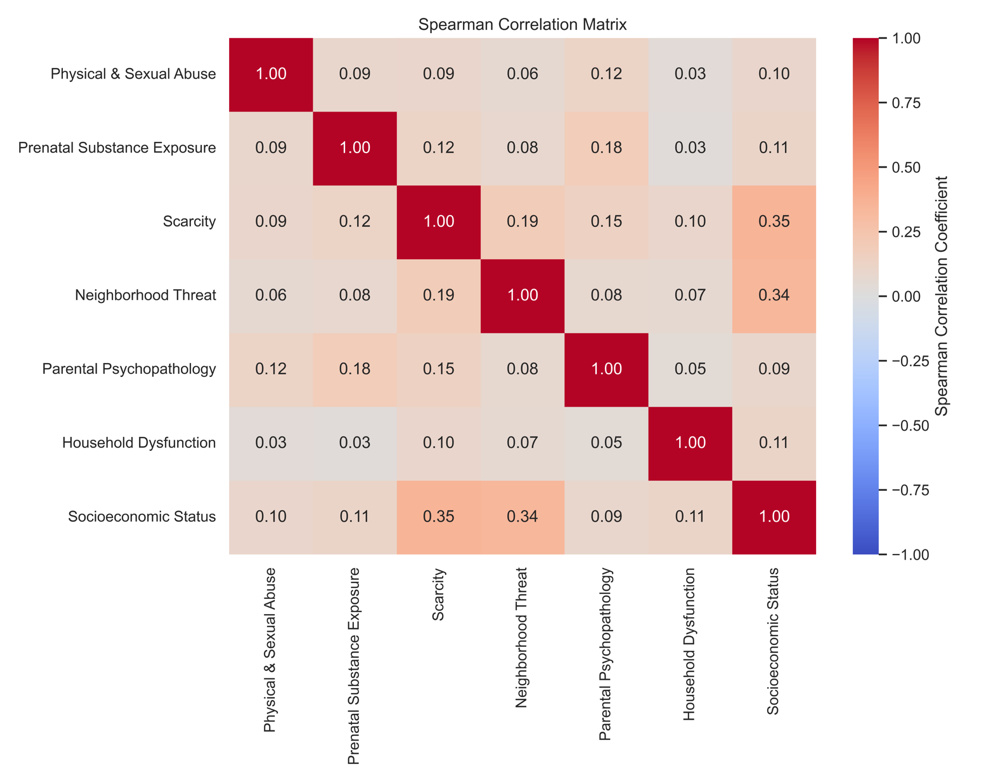

**Supplementary Figure 2: Behavioral measures are weakly to moderately correlated at baseline.**

(A) Histograms describe distribution of adversity factors and socioeconomic status for participants at baseline. Physical and Sexual Abuse range 0-9, Neighborhood Threat range 0-8, Prenatal Substance Exposure range 0-6, Household Dysfunction 0-3, Scarcity range 0-2, Parental Psychopathology range 0-16, Household Dysfunction 0-3, Socioeconomic Status (measured using household income) range 1-10. Higher scores indicate higher adversity/lower income.

(B) The 2D grids display the correlation coefficient for each pair of behavioral scores.

**Participant Inclusion Pipeline at Two-Year Follow-Up**

*n* = 10,908

*n* = 3 excluded: intersex individuals

n = 10,905

*n* = 2,825 excluded: missing brain structural data

*n* = 8,080

*n* = 7,884

*n* = 196 excluded: T1 data not recommended for inclusion, (“imgincl_t1w_include = 0”)

*n* = 357 excluded: T1 data incidental findings reported (“mrif_score = 3 or 4”)

*n* = 7,527

*n* = 6,781

*n* = 5,819

***n* = 5,808**

*n* = 11 excluded: site22 < 20 individuals, (“site_id < 20”)

*n* = 1,103 excluded: missing behavioral data to calculate adversity factors or income

*n* = 962 excluded: siblings, (duplicate “rel_family_id”)

**Supplementary Figure 3: Participant inclusion and exclusion workflow at two-year follow-up.**

Overview of the process used to include participants in the analyses at two-year follow-up.

A. **Distribution of Behavioral Scores at Two-Year Follow-Up**

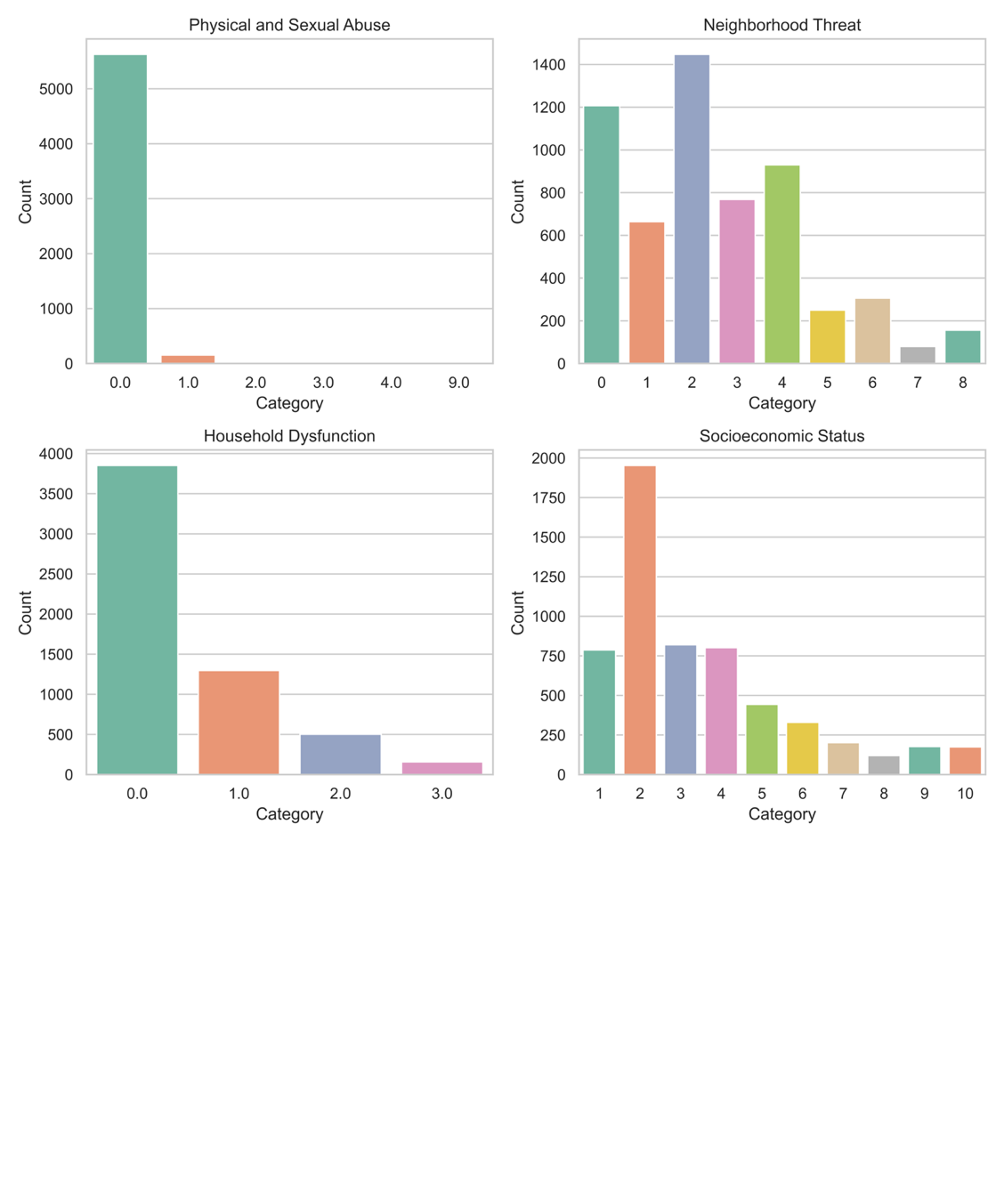

B. **Correlation of Behavioral Scores at Two-Year Follow-Up**

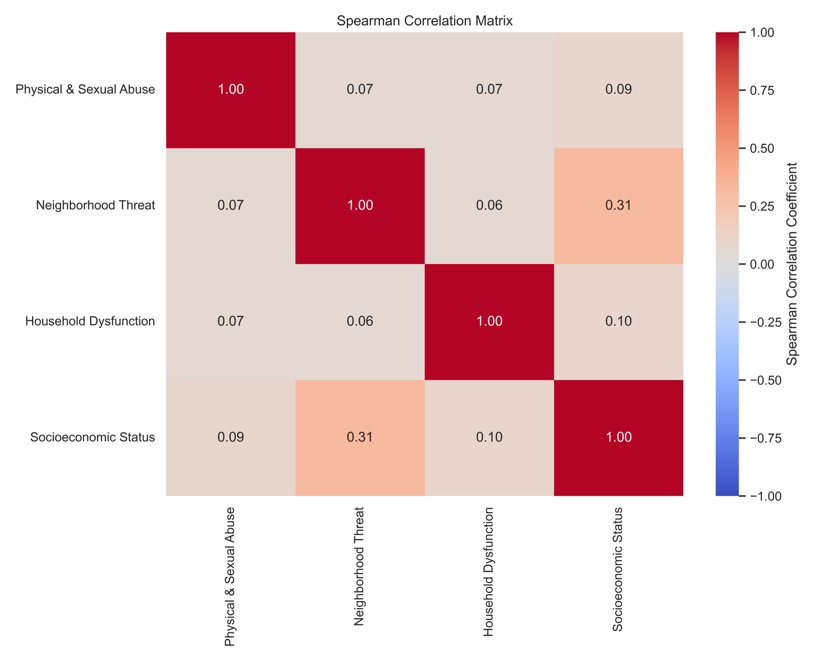

**Supplementary Figure 4: Behavioral measures are weakly to moderately correlated at two-year follow-up**

(A) Histograms describe distribution of adversity factors and socioeconomic status for participants at Two-Year Follow-Up. Physical and Sexual Abuse range 0-9, Neighborhood Threat range 0-8, Household Dysfunction 0-3, Socioeconomic Status (measured using household income) range 1-10. Higher scores indicate higher adversity/lower income.

(B) The 2D grids display the correlation coefficient for each pair of behavioral scores.

**Participant Inclusion Pipeline at Four-Year Follow-Up**

*n* = 4,688

*n* = 1 excluded: intersex individual

n = 4,687

*n* = 1,675 excluded: missing brain structural data

*n* = 3,012

*n* = 2,980

*n* = 32 excluded: T1 data not recommended for inclusion, (“imgincl_t1w_include = 0”)

*n* = 167 excluded: T1 data incidental findings reported, (“mrif_score = 3 or 4”)

*n* = 2,813

*n* = 2,583

*n* = 2,267

***n* = 2,245**

*n* = 230 excluded: missing behavioral data to calculate adversity factors or income

*n* = 316 excluded: siblings, (duplicate “rel_family_id”)

*n* = 22 excluded: site22, site7, < 20 individuals. (“site_id < 20”)

**Supplementary Figure 5: Participant inclusion and exclusion workflow at four-year follow-up.**

Overview of the process used to include participants in the analyses at four-year follow-up.

A. **Distribution of Behavioral Scores at Four-Year Follow-Up**

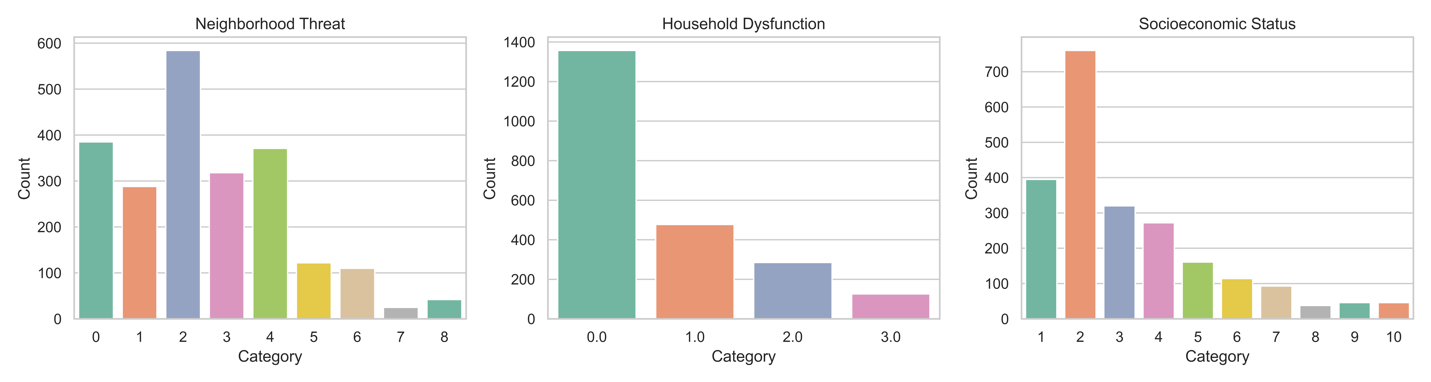

B. **Correlation of Behavioral Scores at Four-Year Follow-Up**

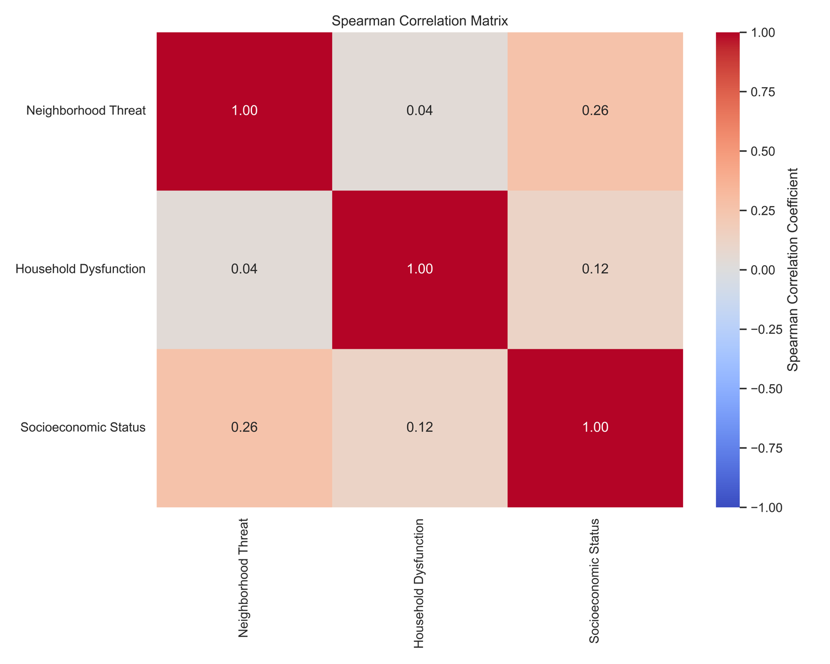

**Supplementary Figure 6: Behavioral measures are weakly to moderately correlated at four-year follow-up**

(A) Histograms describe distribution of adversity factors and socioeconomic status for participants at Four-Year Follow-Up. Neighborhood Threat range 0-8, Household Dysfunction 0-3, Socioeconomic Status (measured using household income) range 1-10. Higher scores indicate higher adversity/lower income.

(B) The 2D grids display the correlation coefficient for each pair of behavioral scores.

**Participant Overlap across time-points**

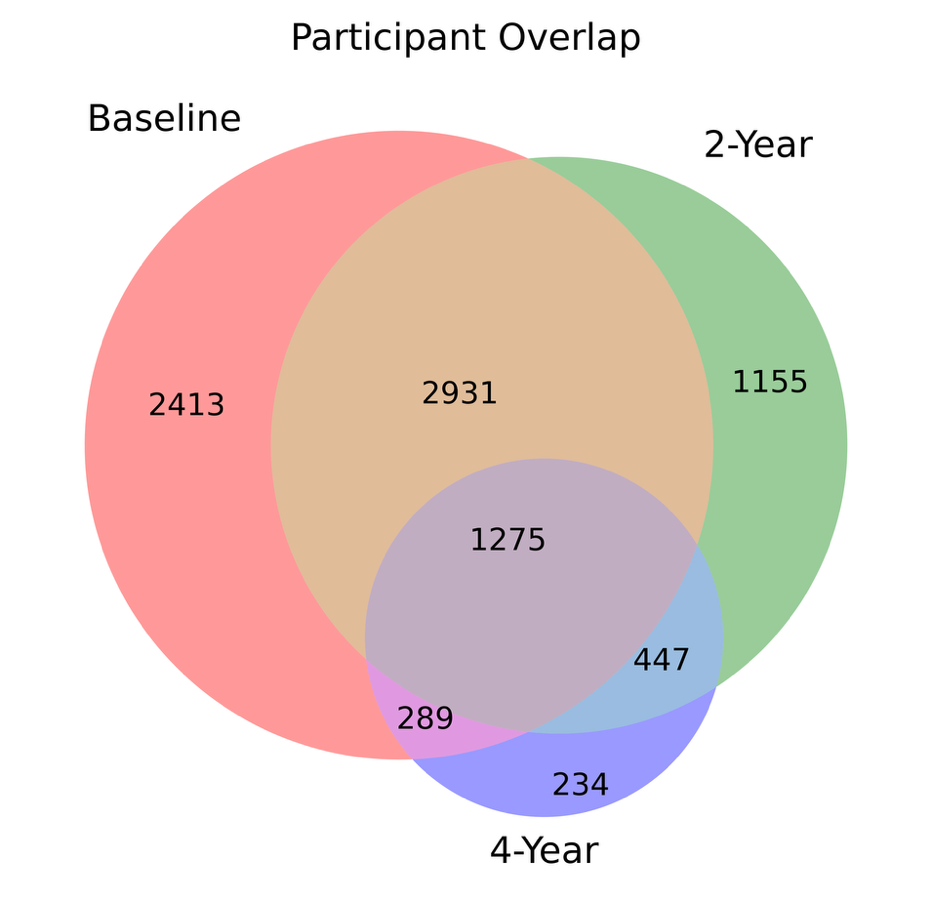

**Supplementary Figure 7: Participant overlap across the three timepoints included in the analysis.** Baseline (red circle), two-year follow-up (green circle), and four-year follow-up (blue circle). Numbers indicate how many participants contributed data at each individual timepoint and at overlapping combinations of timepoints.

**Relative associations between cortical thickness and significant factors for sex-specific analyses at baseline**

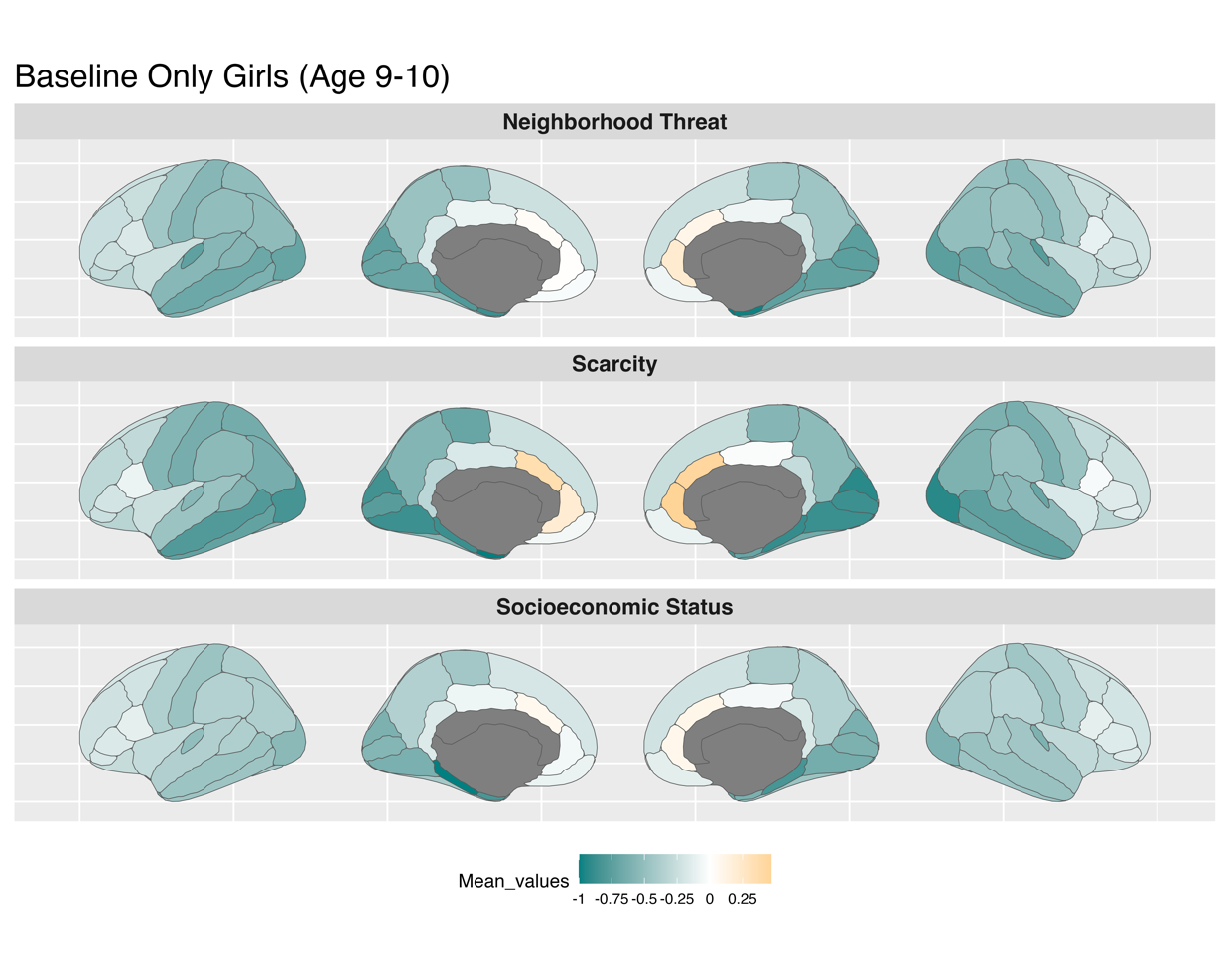

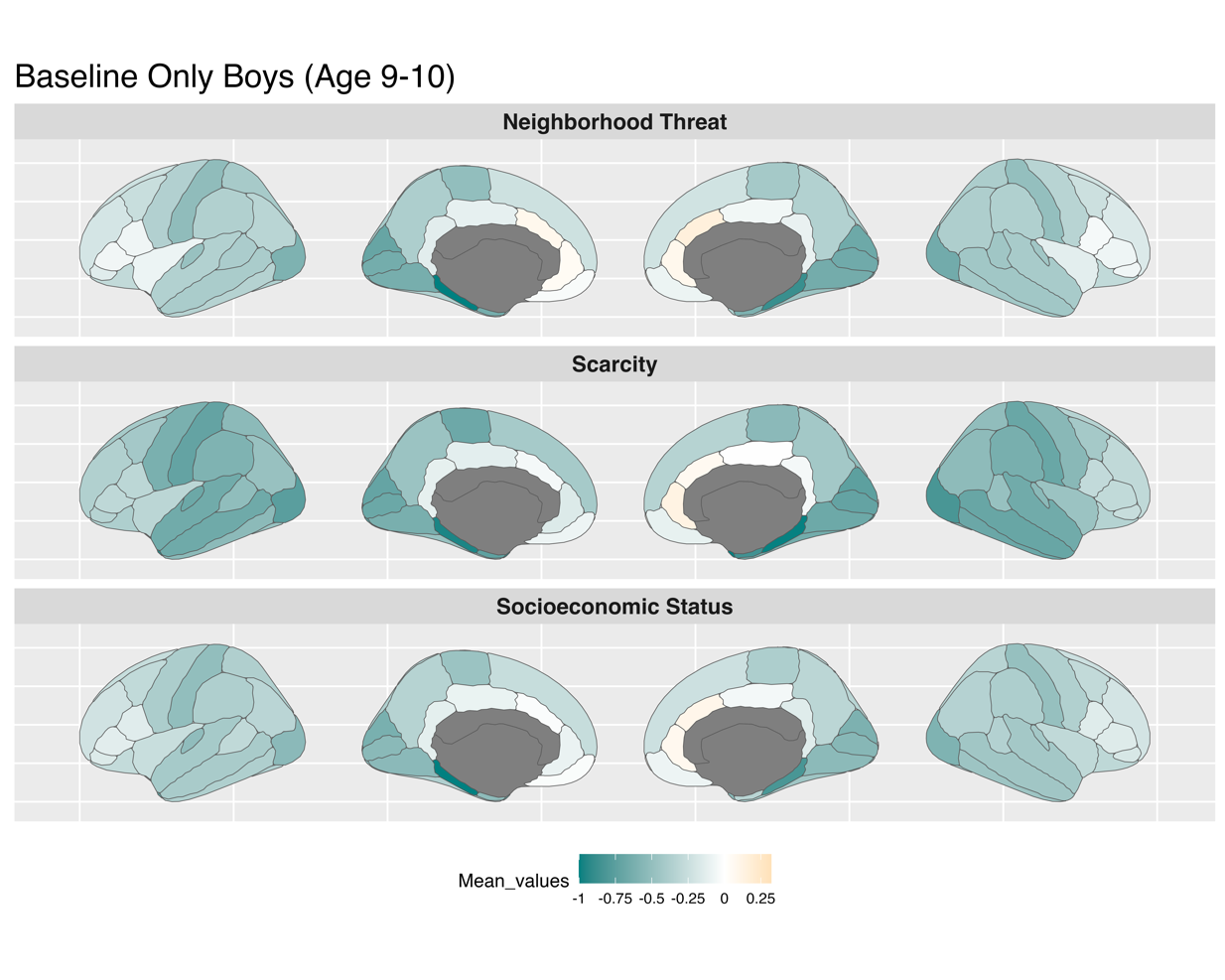

**Supplementary Figure 8: Baseline - Feature Weights for Sex-specific Models**

**Relative associations between cortical thickness and significant factors for sex-specific analyses at two-year follow-up**

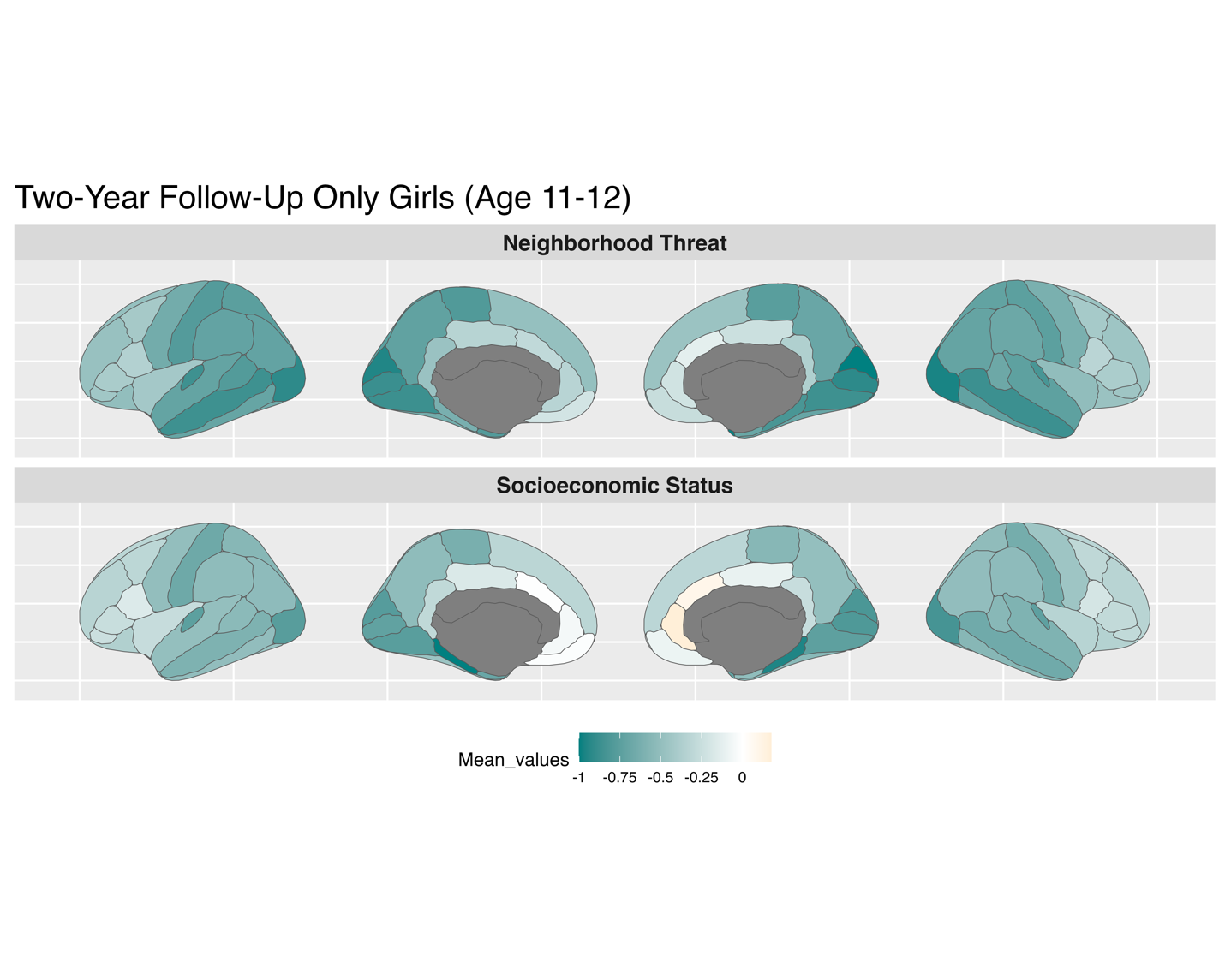

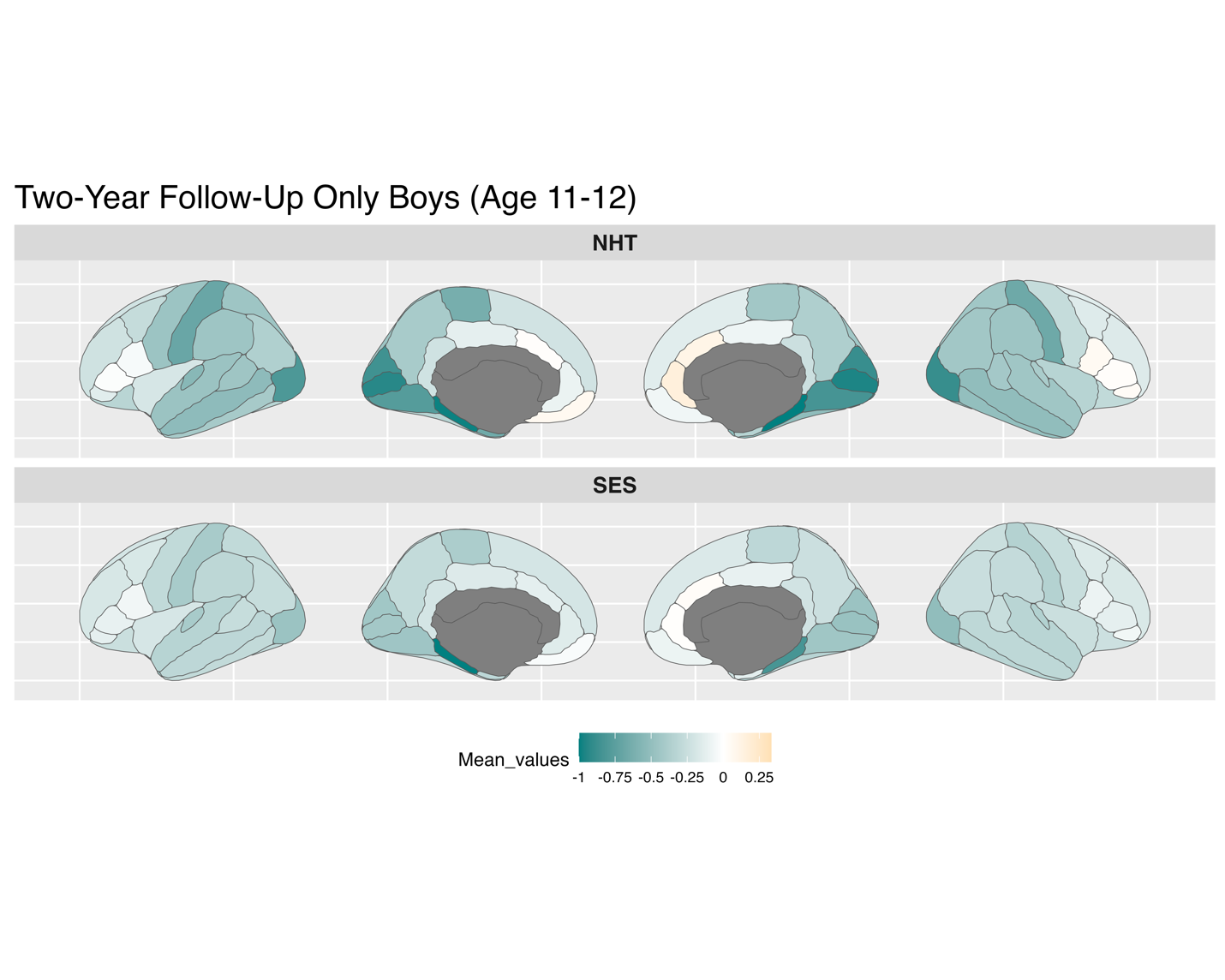

**Supplementary Figure 9: Two-Year Follow-Up Feature Weights for Sex-specific Models**

**Relative associations between cortical thickness and significant factors for sex-specific analyses at four-year follow-up**

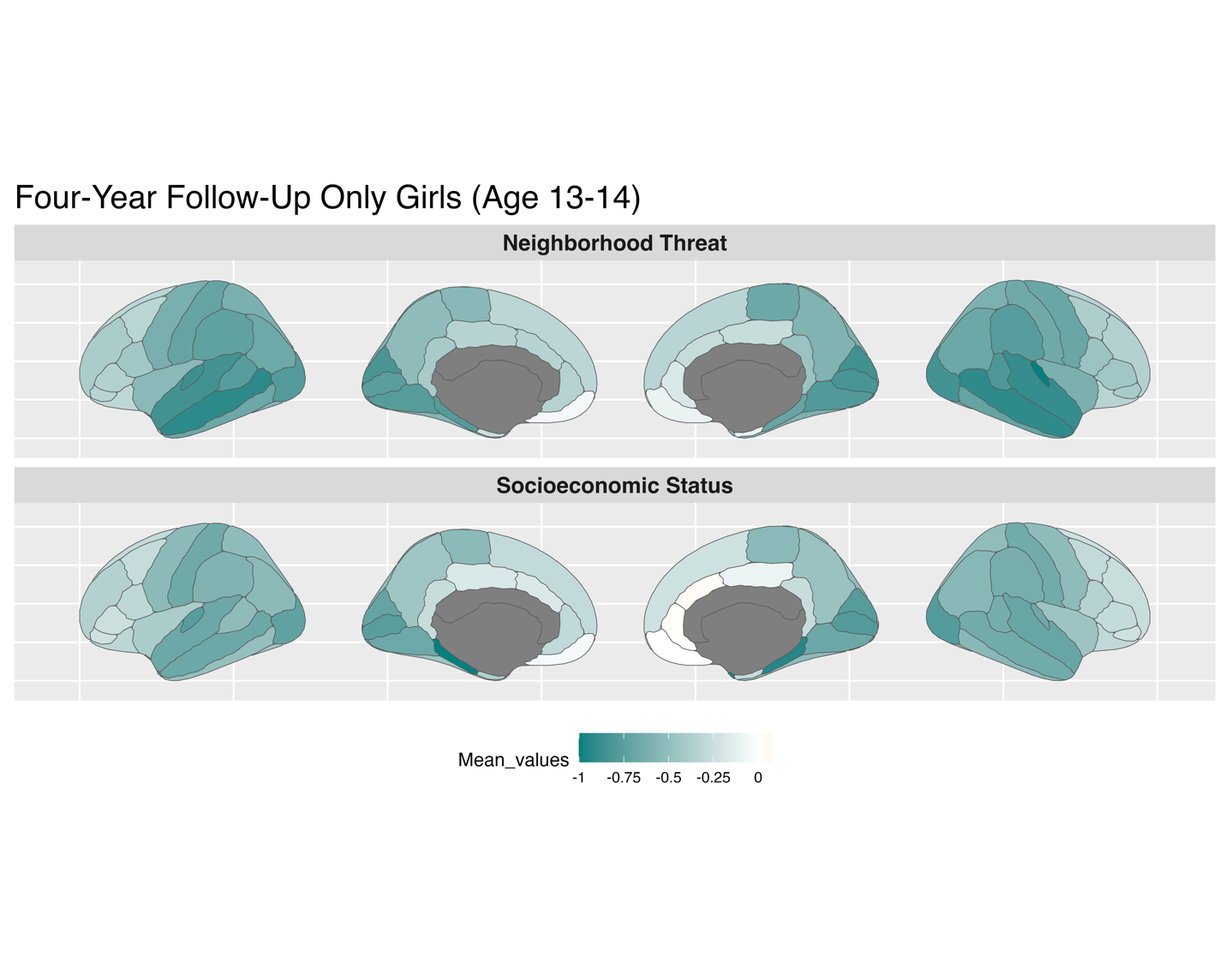

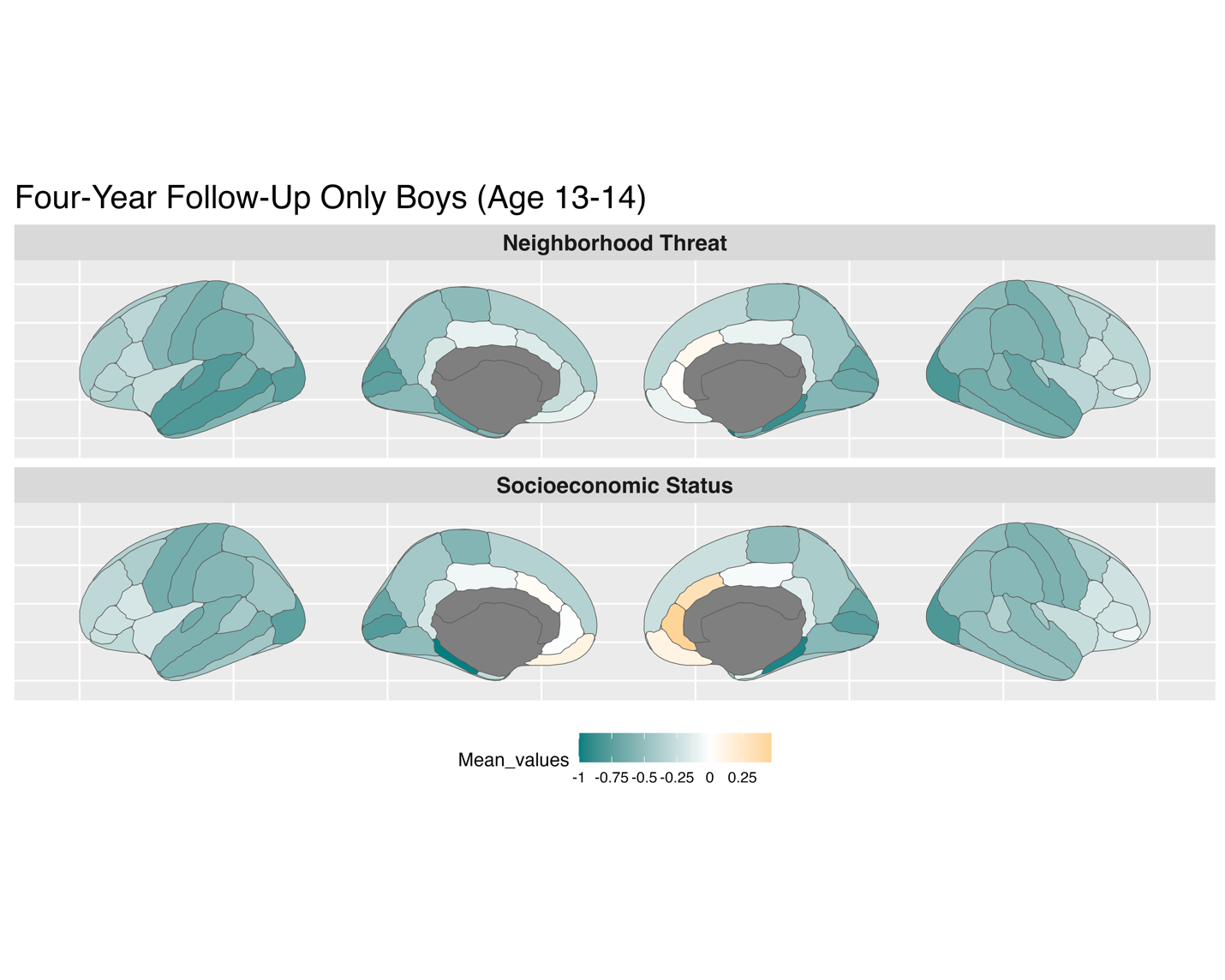

**Supplementary Figure 10: Four-Year Follow-Up - Feature Weights for Sex-specific Models**

**Cosine Similarity at Baseline, Two-Year and Four-Year Follow-Up**

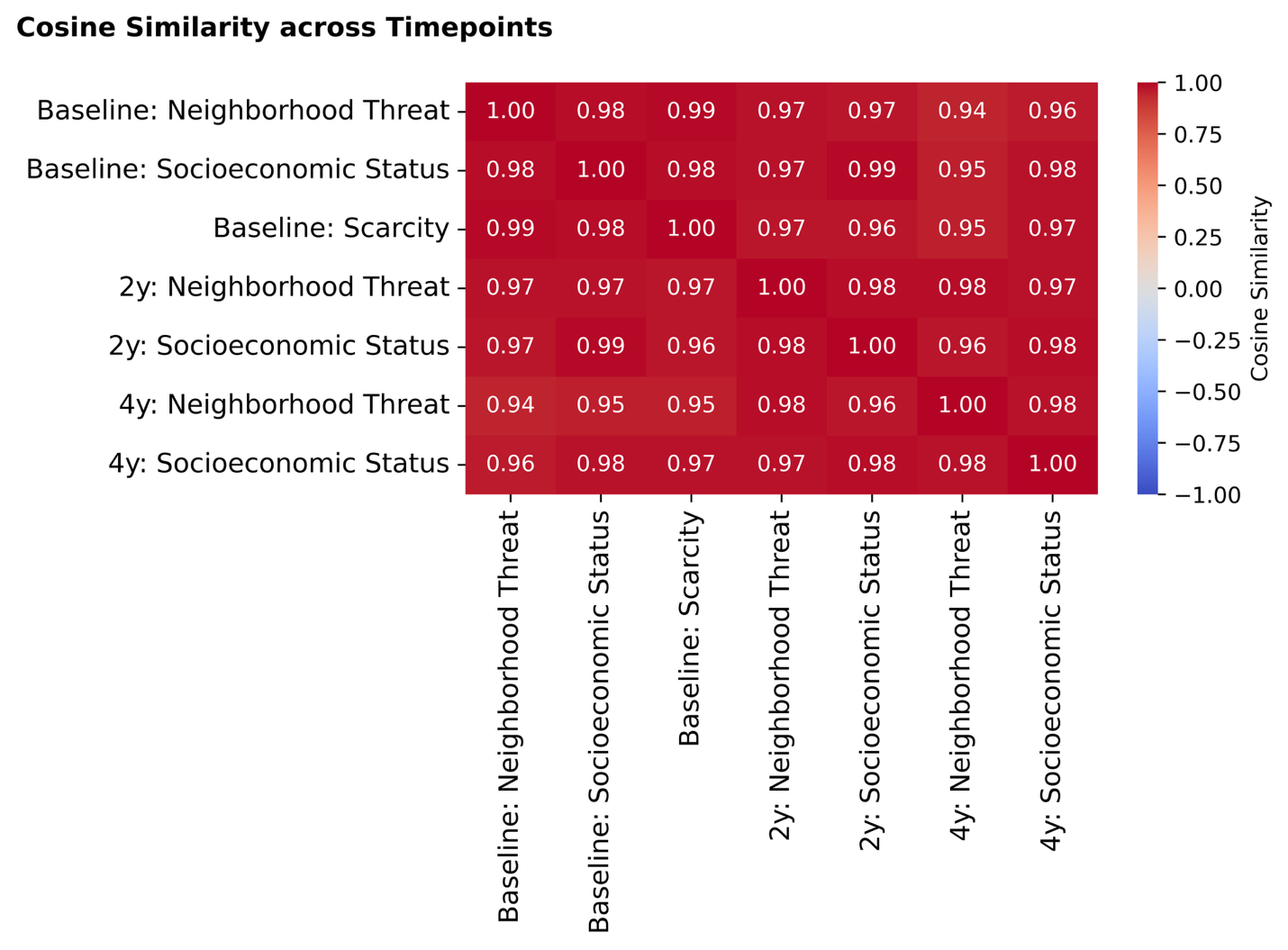

**Supplementary Figure 11: Cosine similarities for feature importance in the significant models across time-points**. Cosine similarities between the Haufe-transformed regional feature weights from models trained to predict early life adversity and socioeconomic status across three time-points. Results for models based that captured significant associations are shown. Warmer colors indicate greater similarity, cooler colors indicate a greater dissimilarity.

**Robustness Analysis: Relative associations between cortical thickness and Income-to-needs Ratio**

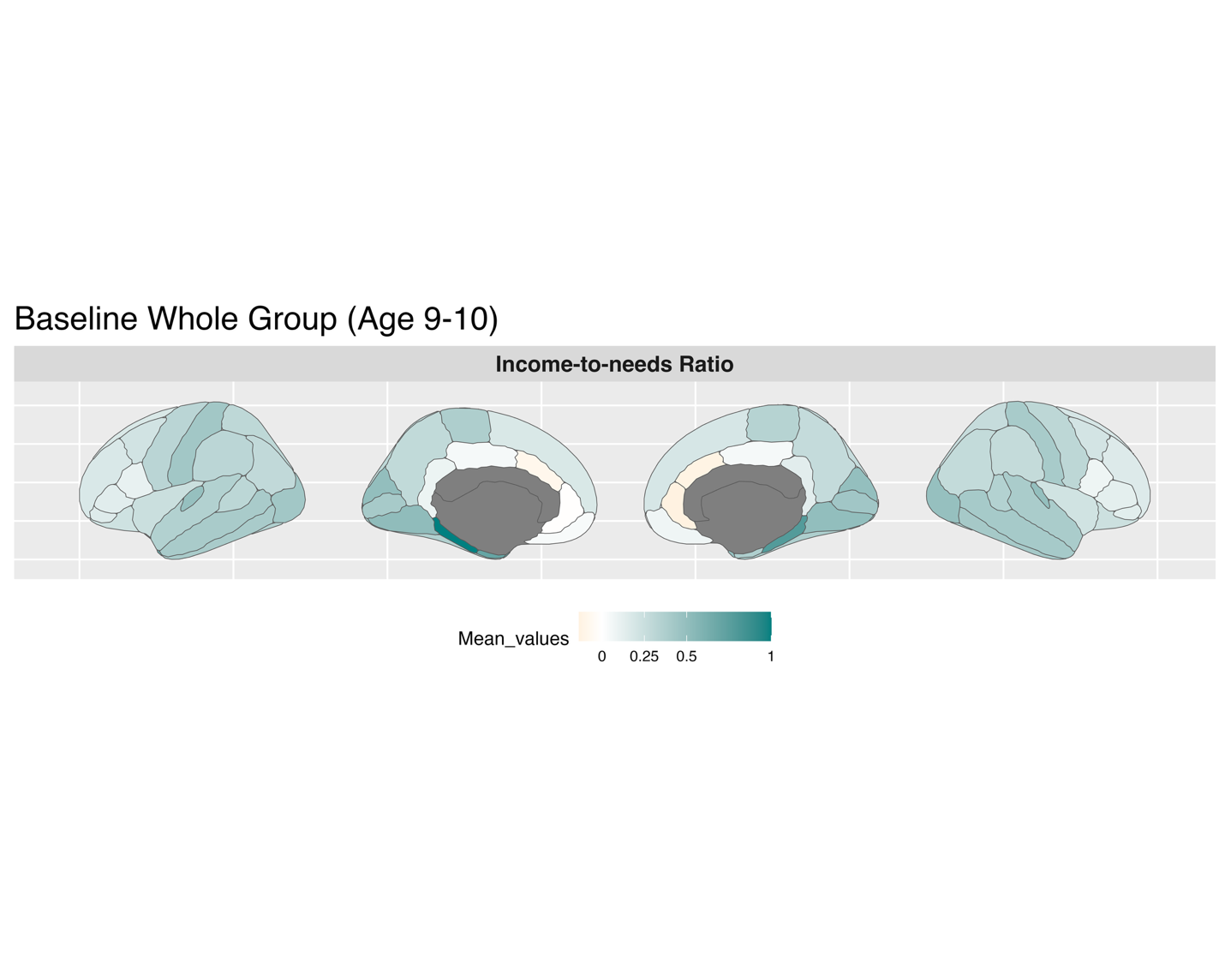

**Supplementary Figure 12: Baseline - Feature Weights for Whole Group for Income-to-needs Ratio.** Income-to-needs ratio is a more sensitive measure of socioeconomic status. At baseline, income-to-needs ratio was significantly associated with cortical thickness (r=0.220, p<0.001).

**Robustness analysis: Sample with zero-overlap in participants across time-points.**

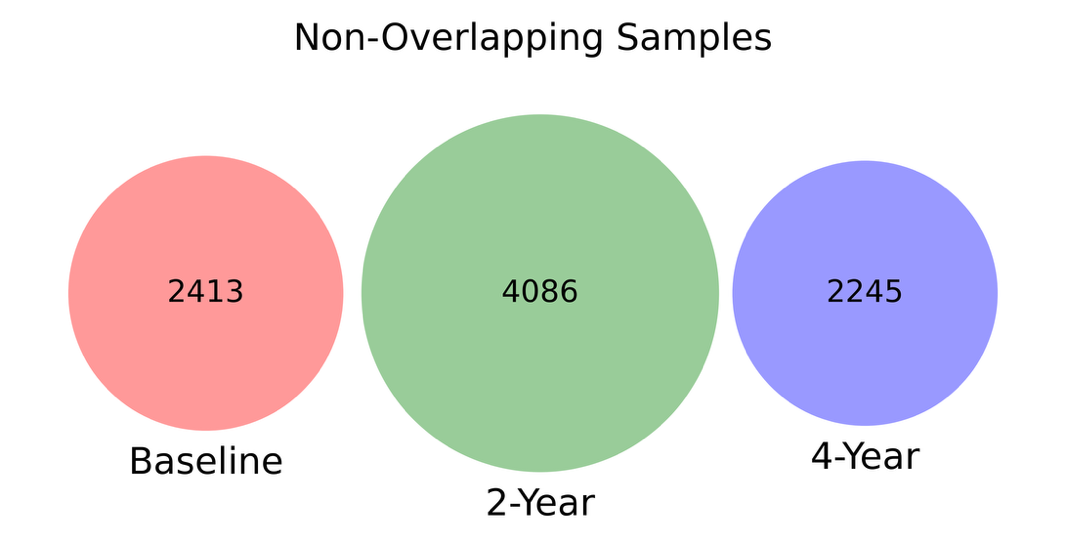

**Supplementary Figure 13: Sample sizes for non-overlapping sample sizes**

Sample sizes for the robustness analysis using non-overlapping participants at each timepoint. Each group includes only unique individuals who contributed data at a single timepoint (baseline, two-year follow-up, or four-year follow-up), ensuring zero overlap across timepoints.

**Robustness Analysis for non-overlapping sample.**

To ensure that our consistent findings were not driven by repeated measurements of the same participants, we ran additional robustness analysis with non-overlapping samples across time-points. To ensure maximum participants at each time-point, we maintained our four year-analysis data, and reran the analysis for two-year follow-up (n= 4086) and baseline (n= 2413), ensuring zero participant overlap across time points.

At baseline, cortical thickness was significantly associated with neighborhood threat (prediction accuracy, r=0.158, pFDR = 0.011), scarcity (r=0.101, pFDR=0.030), and socioeconomic status (r=0.275, pFDR<0.001). Cortical thickness was not significantly associated with physical and sexual abuse (r=-0.009, pFDR=0.561), prenatal substance exposure (r=0.011, pFDR=0.462), household dysfunction (r=0.026, pFDR=0.440), or parental psychopathology (r=0.018, pFDR=0.440), indicating similar findings as in the main analysis.

At two-year follow-up, cortical thickness was significantly associated with neighborhood threat (r=0.114, pFDR = 0.006) and socioeconomic status (r=0.280, pFDR<0.001). Cortical thickness was not significantly associated with physical and sexual abuse (r=0.009, pFDR=0.322) or household dysfunction (r=0.031, pFDR=0.157), indicating similar findings as in the main analysis.

**Robustness Analysis: Relative associations between cortical thickness and significant factors for zero-overlap samples at baseline and two-year follow-up.**

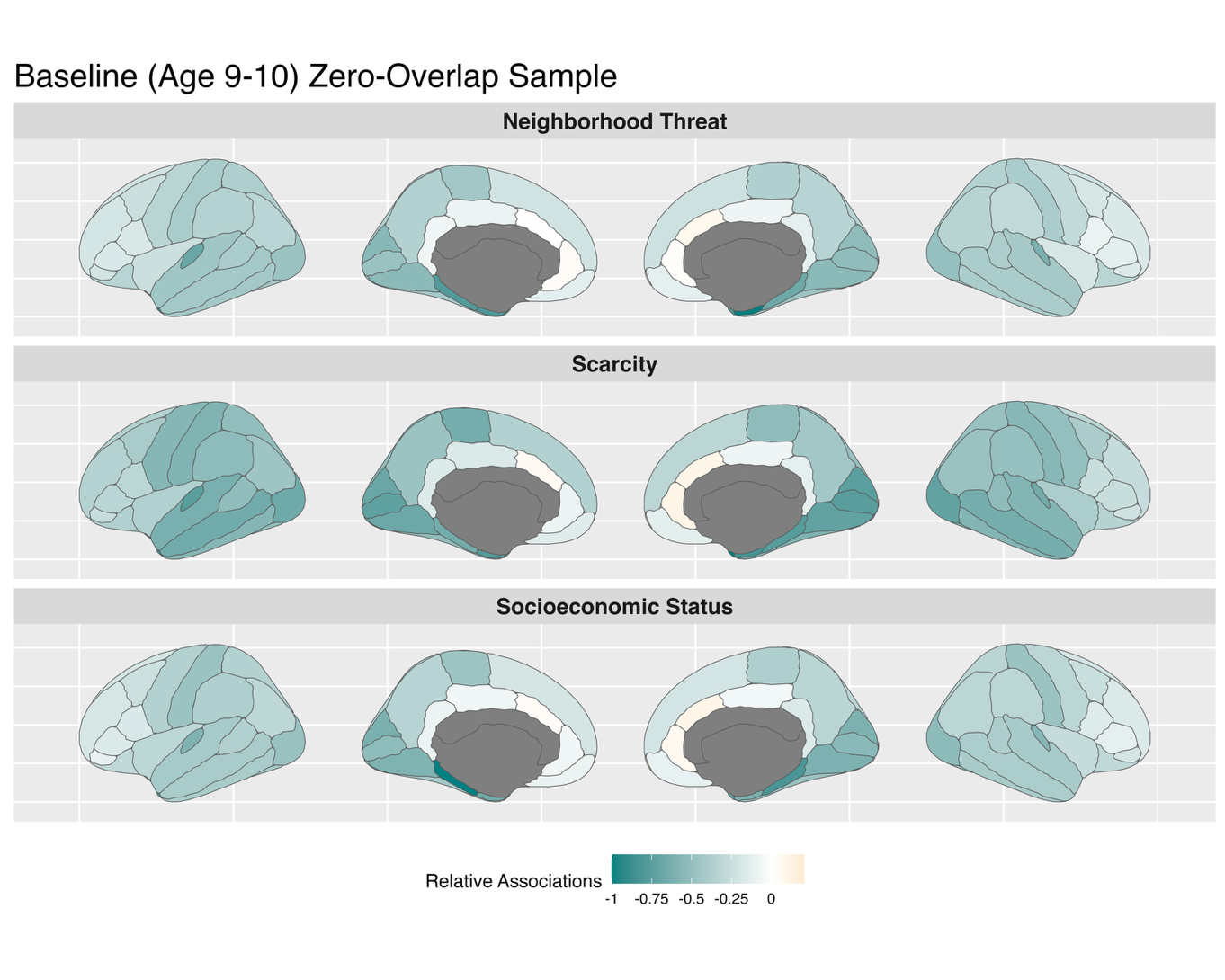

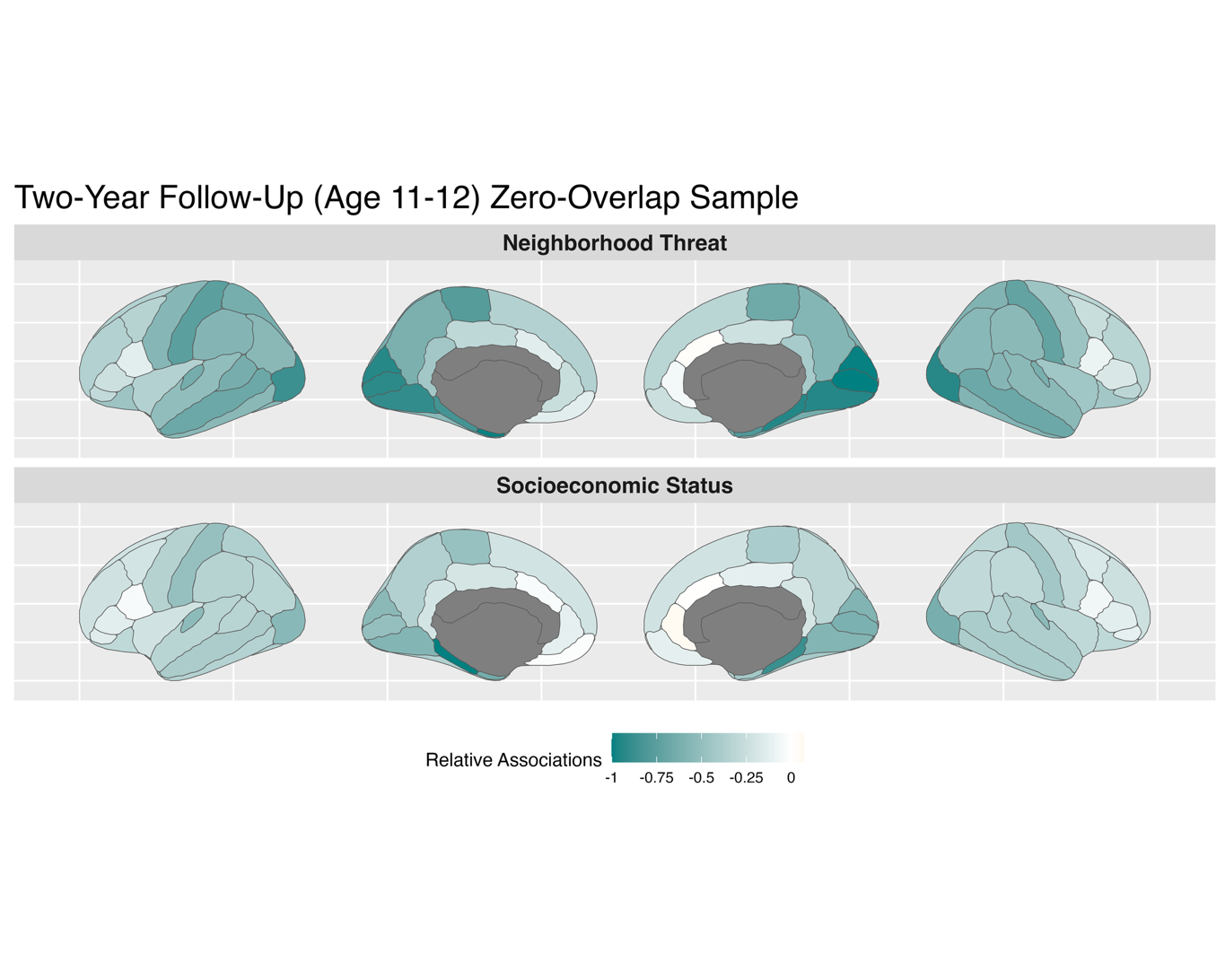

**Supplementary Figure 14: Feature weights for the non-overlapping samples at baseline and two-year follow-up.** Findings remain robust in completely independent samples across time-points.

**Changes in adversity exposure over time.**

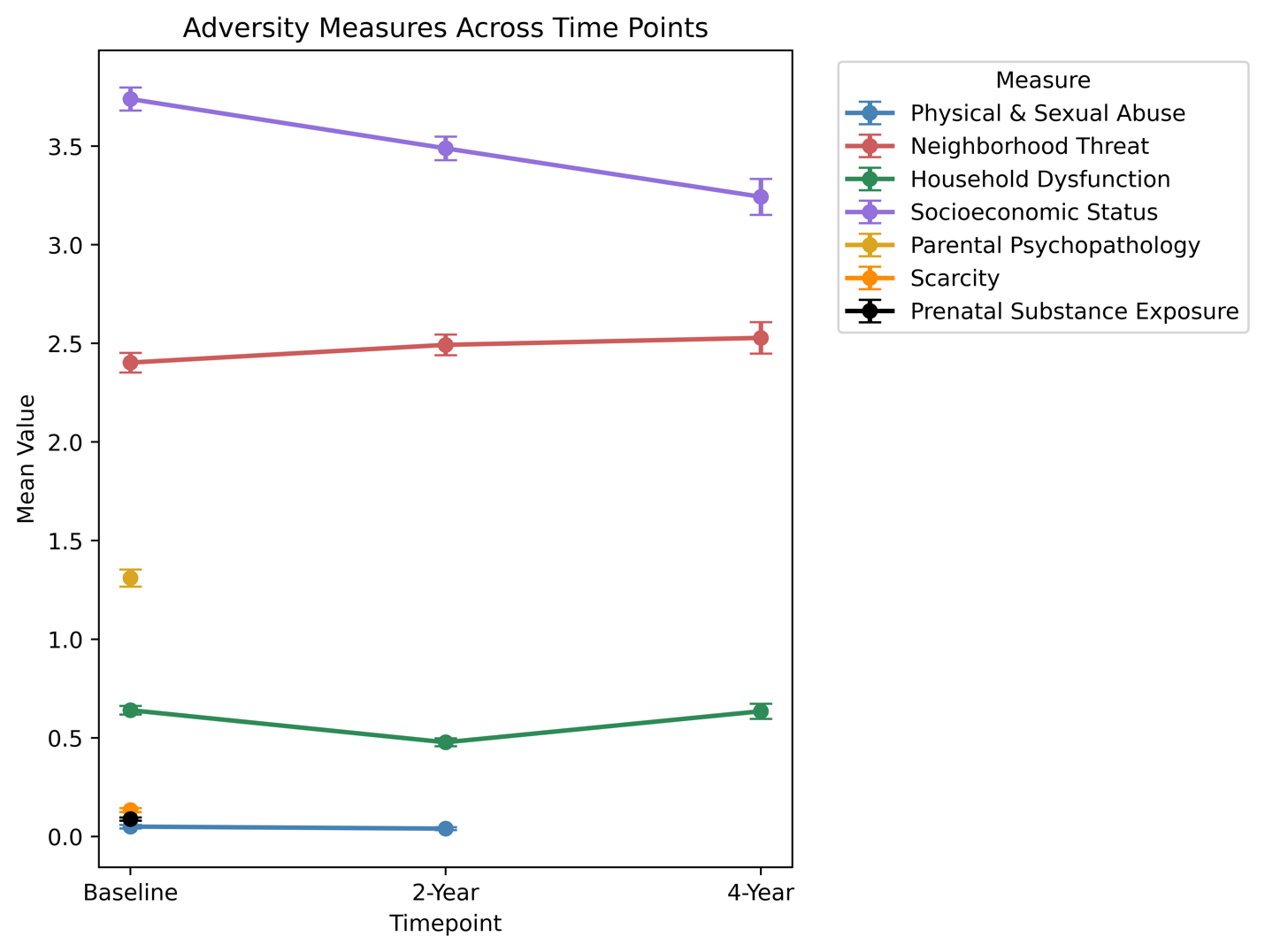

**Supplementary Figure 15: Behavioral measures remain relatively stable across time.**

Dots indicate mean, Error bars indicate two standard error of the mean (SEM). Scarcity, Parental Psychopathology, and Prenatal Substance Exposure measures were only available at baseline. Physical and Sexual Abuse measure was only available at baseline and two-year follow-up.

**Calculation of early life adversity scores according to factor analysis by Orendain et al., 2023**

**Physical and Sexual Violence**

1. Shot, stabbed, or beaten brutally by a non-family member? From: Kiddie Schedule for Affective Disorders and Schizophrenia (KSADS-5)
2. Shot, stabbed, or beaten brutally by a grown up in the home? From: Kiddie Schedule for Affective Disorders and Schizophrenia (KSADS-5)
3. Beaten to the point of having bruises by a grown up in the home? From: Kiddie Schedule for Affective Disorders and Schizophrenia (KSADS-5)
4. A grown up in the home touched your child in his or her privates, had your child touch their privates, or did other sexual things to your child? From: Kiddie Schedule for Affective Disorders and Schizophrenia (KSADS-5)
5. An adult outside your family touched your child in his or her privates, had your child touch their privates, or did other sexual things to your child? From: Kiddie Schedule for Affective Disorders and Schizophrenia (KSADS-5)
6. A peer forced your child to do something sexually? From: Kiddie Schedule for Affective Disorders and Schizophrenia (KSADS-5)
7. Witnessed someone shot or stabbed in the community From: Kiddie Schedule for Affective Disorders and Schizophrenia (KSADS-5)
8. A non-family member threatened to kill your child? From: Kiddie Schedule for Affective Disorders and Schizophrenia (KSADS-5)
9. A family member threatened to kill your child? From: Kiddie Schedule for Affective Disorders and Schizophrenia (KSADS-5)

**Parental Psychopathology**

1. Has the child’s biological parents ever had any problems due to alcohol, such as: Marital separation or divorce; Laid off or fired from work; Arrests or DUIs; Alcohol harmed their health; In an alcohol treatment program; Suspended or expelled from school 2 or more times; Isolated self from family, caused arguments or were drunk a lot. From: ABCD Family History Assessment
2. Has the child’s biological parents ever had any problems due to drugs, such as: Marital separation or divorce; Laid off or fired from work; Arrests or DUIs; Drugs harmed their health; In a drug treatment program; Suspended or expelled from school 2 or more times; Isolated self from family, caused arguments or were high a lot. From: ABCD Family History Assessment
3. Has the child’s biological parents ever suffered from depression, that is, have they felt so low for a period of at least two weeks that they hardly ate or slept or couldn't work or do whatever they usually do? From: ABCD Family History Assessment
4. Has the child’s biological parents ever had a period of time when others were concerned because they suddenly became more active day and night and seemed not to need any sleep and talked much more than usual for them? From: ABCD Family History Assessment
5. Has the child’s biological parents ever had a period lasting six months when they saw visions or heard voices or thought people were spying on them or plotting against them? From: ABCD Family History Assessment
6. Has the child’s biological parents ever been to a doctor or a counselor about any emotional or mental problems, or problems with alcohol or drugs? From: ABCD Family History Assessment
7. Has the child’s biological parents ever been hospitalized because of emotional or mental problems, or drug or alcohol problems? From: ABCD Family History Assessment
8. Has the child’s biological parents ever attempted or committed suicide? From: ABCD Family History Assessment

**Neighborhood Threat**

1. My neighborhood is safe from crime. From: ABCD Parent Neighborhood Safety/Crime Survey modified from PhenX (NSC)
2. Violence is not a problem in my neighborhood. From: ABCD Parent Neighborhood Safety/Crime Survey modified from PhenX (NSC)

**Prenatal Substance Exposure**

Once you knew you were pregnant, were you using any of the following:

1. tobacco
2. alcohol
3. marijuana
4. cocaine/crack
5. heroin/morphine
6. OxyContin

All From: ABCD Developmental History Questionnaire

**Scarcity**

In the past 12 months, has there been a time when you and your immediate family experienced any of the following:

1. Needed food but couldn’t afford to buy it or couldn’t afford to go out to get it? From: ABCD Parent Demographics Survey
2. Had services turned off by the gas or electric company, or the oil company wouldn’t deliver oil because payments were not made? From: ABCD Parent Demographics Survey

**Household Dysfunction**

1. Family members sometimes hit each other. (Youth-reported) From: ABCD Youth Family Environment Scale-Family Conflict Subscale modified from PhenX (FES)
2. We fight a lot in our family. (Youth-reported) From: ABCD Youth Family Environment Scale-Family Conflict Subscale modified from PhenX (FES)
3. Family members often criticize each other. (Youth-reported) From: ABCD Youth Family Environment Scale-Family Conflict Subscale modified from PhenX (FES)
